# A predictive model of gene expression reveals the role of regulatory motifs in the mating response of yeast

**DOI:** 10.1101/2020.06.23.167205

**Authors:** Amy E. Pomeroy, Matthew I. Pena, John R. Houser, Gauri Dixit, Henrik G. Dohlman, Timothy C. Elston, Beverly Errede

## Abstract

Cells use signaling pathways to receive and process information about their environment. These systems are nonlinear, relying on feedback and feedforward regulation to respond appropriately to changing environmental conditions. Mathematical models developed to describe signaling pathways often fail to show predictive power, because the models are not trained on data that probe the diverse time scales on which feedforward and feedback regulation operate. We addressed this limitation using microfluidics to expose cells to a broad range of dynamic environmental conditions. In particular, we focus on the well-characterized mating response pathway of *S. cerevisiae* (yeast). This pathway is activated by mating pheromone and initiates the transcriptional changes required for mating. Although much is known about the molecular components of the mating response pathway, less is known about how these components function as a dynamical system. Our experimental data revealed that pheromone-induced transcription persists following removal of pheromone and that long-term adaptation of the transcriptional response occurs when pheromone exposure is sustained. We developed a model of the regulatory network that captured both persistence and long-term adaptation of the mating response. We fit this model to experimental data using an evolutionary algorithm and used the parameterized model to predict scenarios for which it was not trained, including different temporal stimulus profiles and genetic perturbations to pathway components. Our model allowed us to establish the role of four regulatory motifs in coordinating pathway response to persistent and dynamic stimulation.

## INTRODUCTION

Proper cellular function requires cells to respond appropriately to stimuli in their environment. Environmental cues, such as hormones and growth factors, are typically sensed by receptors on the cell surface and transmitted by intracellular signaling pathways. A key function of these pathways is to initiate the appropriate transcriptional program to respond to the environmental challenge. Mathematical modeling has helped to elucidate many of the design principles that regulate the spatiotemporal activity of signaling pathways and allow them to function reliably in changing environmental conditions *(1)*. The ultimate test for these models is to predict pathway dynamics under conditions of time-dependent stimulation regimens and in the presence of genetic or pharmacological perturbations that disrupt the system in well-defined ways. While many models have reproduced qualitative features of signaling systems, their quantitative predictive power is often lacking. One reason for the lack of predictive power is that many previous studies have assessed cellular responses only to constant stimuli. However, signaling networks are nonlinear systems which typically have both positive and negative feedforward and feedback loops that operate on different time scales. Therefore, full characterization of these systems requires using time-dependent stimulus profiles that probe multiple time scales *(2–12)*.

We have performed such an analysis using the mating response of *Saccharomyces cerevisiae* (yeast). This response is activated when a mating-type specific pheromone binds to and activates a G-protein coupled receptor on a cell of opposite mating type. The signal is then propagated by a mitogen activated protein kinase (MAPK) cascade (**Fig. 1**). A key function of the terminal kinases in this cascade, Fus3 and Kss1, is to initiate the transcriptional program required for successful mating by promoting dissociation of the transcriptional repressors Dig1 and Dig2 from the transcription factor, Ste12 *(13–18)*. Additionally, Fus3 activates Far1, a protein required for cell cycle arrest *(19–21)*. Far1 is also known to affect the transcriptional response by promoting degradation of Ste12 *(22)*. This signaling pathway provides an ideal model system for studying signal transduction and transcriptional regulation *(23, 24)* and has long served as a prototype for MAPK pathways *(25)*. It achieved this status because of the unparalleled ease of genetic manipulation of individual components and unambiguous determination of how these perturbations affect *in vivo* processes. In eukaryotic cells, MAPKs mediate responses to growth factors, cytokines, hormones, cell adhesion, stress and nutrients that determine a wide range of cellular decision processes *(26)*. Thus, a systems level analysis of the yeast mating response is likely to reveal properties common to MAPK regulation of these wide-ranging responses in other cells.

**Figure 1.**
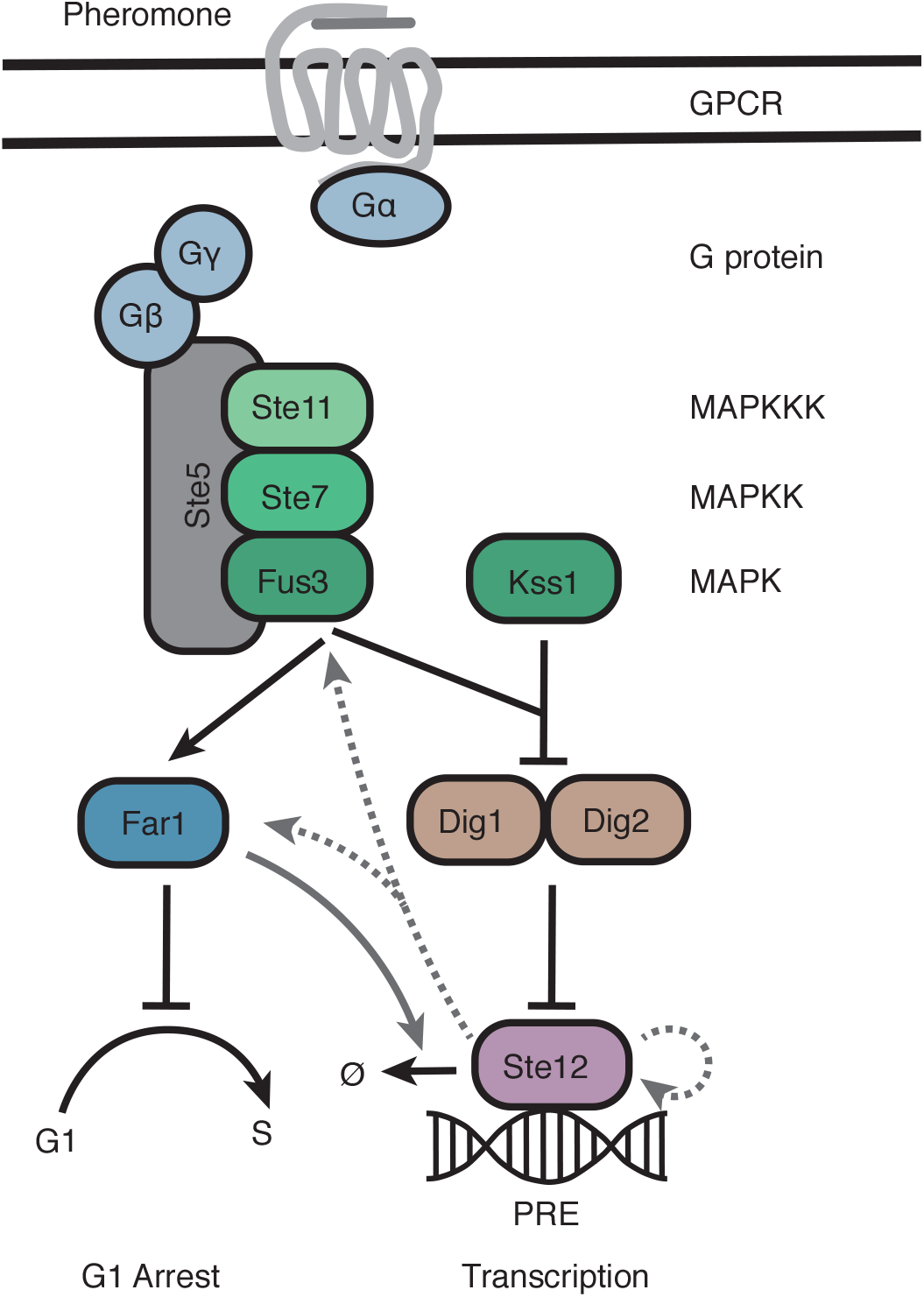
Schematic of the yeast mating response. The yeast mating response is activated when mating pheromone binds to the G-protein coupled receptor (GPCR) activating a heterotrimeric G-protein. The G-protein βγ dimer then activates a mitogen activated protein kinase (MAPK) cascade, resulting in activation of two MAPKs, Fus3 and Kss1. Both kinases activate the transcription factor Ste12 by suppressing the activity of the transcriptional repressors Dig1 and Dig2. Fus3 also activates Far1, which is responsible for cell cycle arrest and promotes degradation of Ste12 (solid gray arrow). Ste12 promotes the transcription of itself, Far1, and Fus3 (dashed gray arrows).

We combined a microfluidics system that allows cells to be exposed to pheromone concentrations with precisely defined temporal profiles and a short-lived fluorescent reporter to monitor dynamic changes in mating specific gene expression. We discovered that transcriptional regulation was sustained following removal of pheromone, a property of the system that we refer to as “persistence”. To better define this surprising property of the system, we exposed cells to pheromone concentrations that oscillate at six different frequencies. The fluorescent data were used to develop and train a model for transcriptional regulation during the mating response. Two strategies were used to validate the model and demonstrate its predictive power. First, we used the model to predict the behavior of mutations that selectively disrupt various signaling motifs in the pathway. Then we used the model to predict the transcriptional response of the system at a lower pheromone concentration. The result of our investigations is a fully validated model of transcriptional regulation that allows a quantitative characterization of the signaling motifs that regulate gene expression. We anticipate that our approach provides a template for a research strategy to characterize regulatory motifs inherent to many signaling pathways.

## RESULTS

### Adaptation and persistence in the mating response pathway

To determine the dynamics of the yeast mating response, we developed experimental tools that allow cells to be exposed to well-defined input signals of any specified temporal profile and a readout that faithfully tracks the dynamic response of the pathway. For controlling stimulus profiles, we employed a microfluidics system that is an adaptation of the “dial-a-wave” system developed by J. Hasty and colleagues *(27)*. For tracking time-dependent changes in pheromone-induced transcription in living cells, we placed a shortlived fluorescent reporter under the control of the pheromone responsive *FUS1* promoter. The fluorescent protein we used is fast maturing (~15 min) and through use of an N-degron tag (YΔk) was engineered to have a half-life similar to its mRNA (~ 7 min) *(28)*. The short-lived reporter is essential in studies of temporal response dynamics, since it reveals transient response characteristics that are otherwise masked by accumulation of a long-lived reporter protein.

Initially, we exposed cells containing our short-lived fluorescence reporter to a constant stimulus of 50 nM pheromone for 10 hrs and monitored reporter fluorescence by imaging of cells in the microfluidic chamber (**Fig. 2A**). For all the experimental results presented in this manuscript, we used cells lacking the protease Bar1 to remove the effect of pheromone degradation *(29, 30)*. We refer to this strain as wildtype hereafter. Under these conditions, the transcriptional response of wildtype cells reaches a maximum amplitude at 220 min, and then decreases for the remainder of the experiment (**Fig. 2B**).

**Figure 2.**
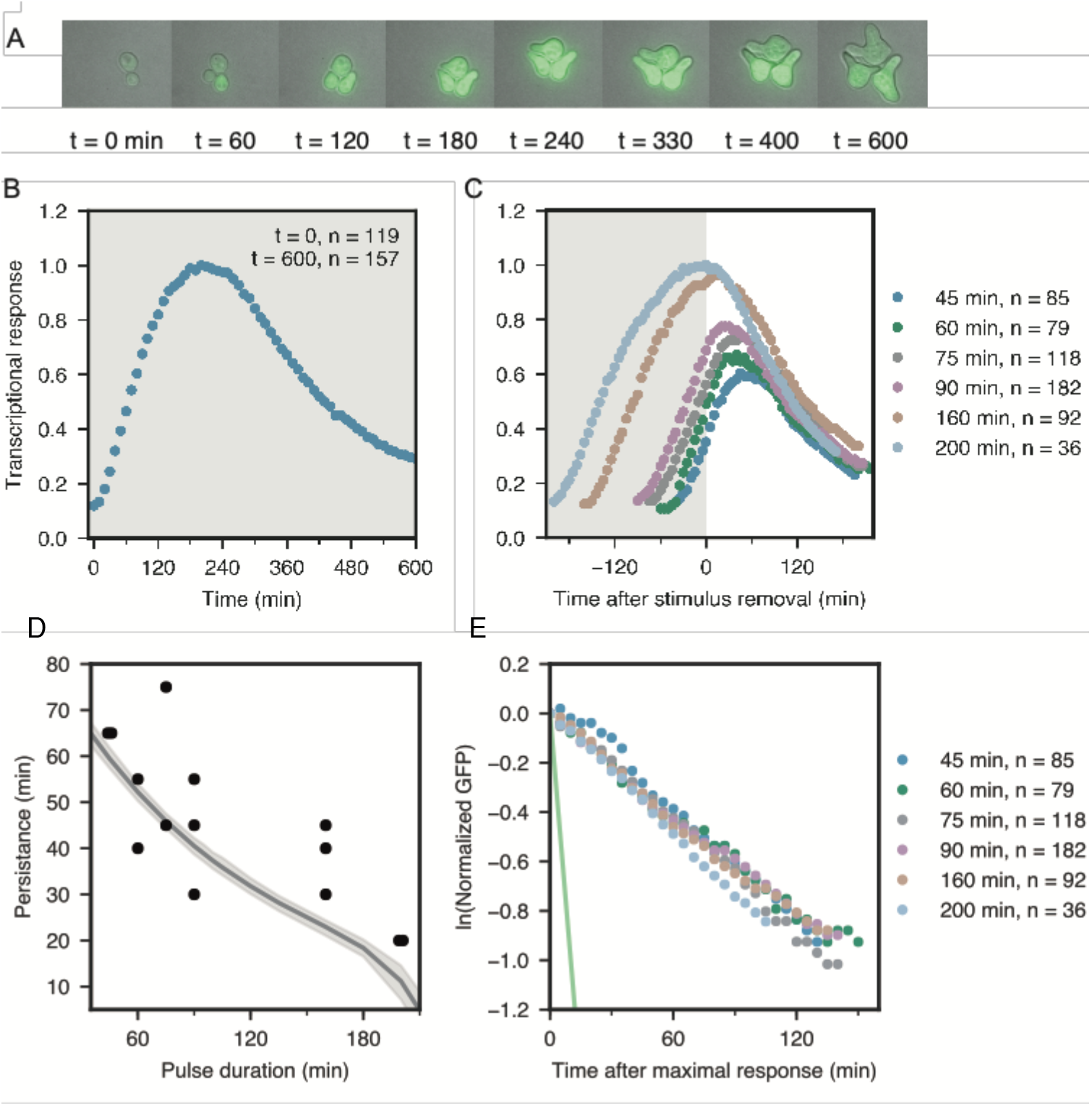
Persistence in transcriptional response. (A) Images of cells with an integrated fluorescent reporter that expresses short-lived GFP from the mating specific *FUS1* promoter exposed to constant stimulus in a microfluidic chamber. (B and C) Quantification of the transcriptional response using fluorescence of the GFP reporter in wildtype (BY4741-68) cells exposed to (B) constant stimulus and (C) stimulus pulses of six different durations (45, 60, 75, 90, 160, and 200 min). (D) The mating transcriptional response persists after a pulse of stimulus is removed, and as the pulse length increases the persistence decreases. This persistence is quantified as the time after stimulus removal until the response to drop 2.5% below the maximum transcription response and each point represents a biological replicate. The solid gray curve represents the mean response from the model and the gray shaded region represents a 99.9% confidence interval band, (E) Adaptation after stimulus removal for each of the six pulse durations is compared by plotting the natural logarithm of the normalized transcriptional response. The normalized response is the average fluorescence at time t after the maximal response divided by the average fluorescence at the onset of adaptation. Assuming exponential decay, the half-life associated with the rate of decreased transcriptional response after stimulus removal is 98 ± 9 min, compared to the 7-minute half-life of the short-lived GFP reporter plotted as a solid green line. Fluorescence data are presented as the average of the indicated number (n) of single cell traces from at least two independent experiments normalized to the maximum response of constant stimulus.

In our next studies, we exposed cells with the short-lived reporter to pheromone pulses of different duration and again monitored reporter fluorescence (**Fig. 2C**). Interestingly, reporter gene expression was significantly sustained following removal of pheromone for pulses of 90 min or less. We refer to this property as persistence and quantify it as the time from removal of pheromone to the time that signal drops below 2.5% of the maximum of each response cuve. The extent of persistence is negatively correlated with the duration of the stimulus pulse; as pulse length increases the persistence of the transcriptional response decreases (**Fig. 2D**). Another important observation is that the rate at which the fluorescent reporter decreases in time is independent of pulse duration (**Fig. 2E**) and the half-life associated with this rate (98 ± 9 min) is considerably longer than the half-lives (~7 min) of the reporter mRNA and protein (**Fig. 2E**, green line) *(28)*. Thus, new synthesis of transcripts and protein continues during the attenuation phase.

A simple explanation for the observed pathway persistence is that it represents a delay between receptor signaling and translation and maturation of the induced GFP reporter. To test this possibility, we developed a linear mathematical model of the response pathway that takes into account this delay (Supplementary Materials). Our analysis of the model revealed that a simple delay cannot account for the persistence in the transcriptional response (**Fig. S1**). In total, our preliminary investigations reveal that the pathway contains some form of “memory” that sustains new mRNA synthesis following removal of pheromone.

We next sought to determine at what level in the pathway the mechanisms for long term adaptation and persistent signaling occur. To determine if “long-term adaptation” relies on upstream pathway regulators of short-term desensitization, such as Sst2 or receptor endocytosis *(31–34)*, we investigated the dynamics of MAPK activity. We monitored MAP kinase dual phosphorylation, which is an indicator of activity, by Western blotting protein extracts of aliquots prepared from cells in the presence of 50 nM pheromone for a 10 hr time course. Fus3 activity remained constant after a transient increase and that of Kss1 increased throughout most of the time course and only slightly diminished toward the end of the experiment (**Fig. 3A**). These results demonstrate that the mechanism of long-term adaptation of transcriptional response does not involve upstream signaling events, but likely occurs at the level of transcriptional regulation.

**Figure 3.**
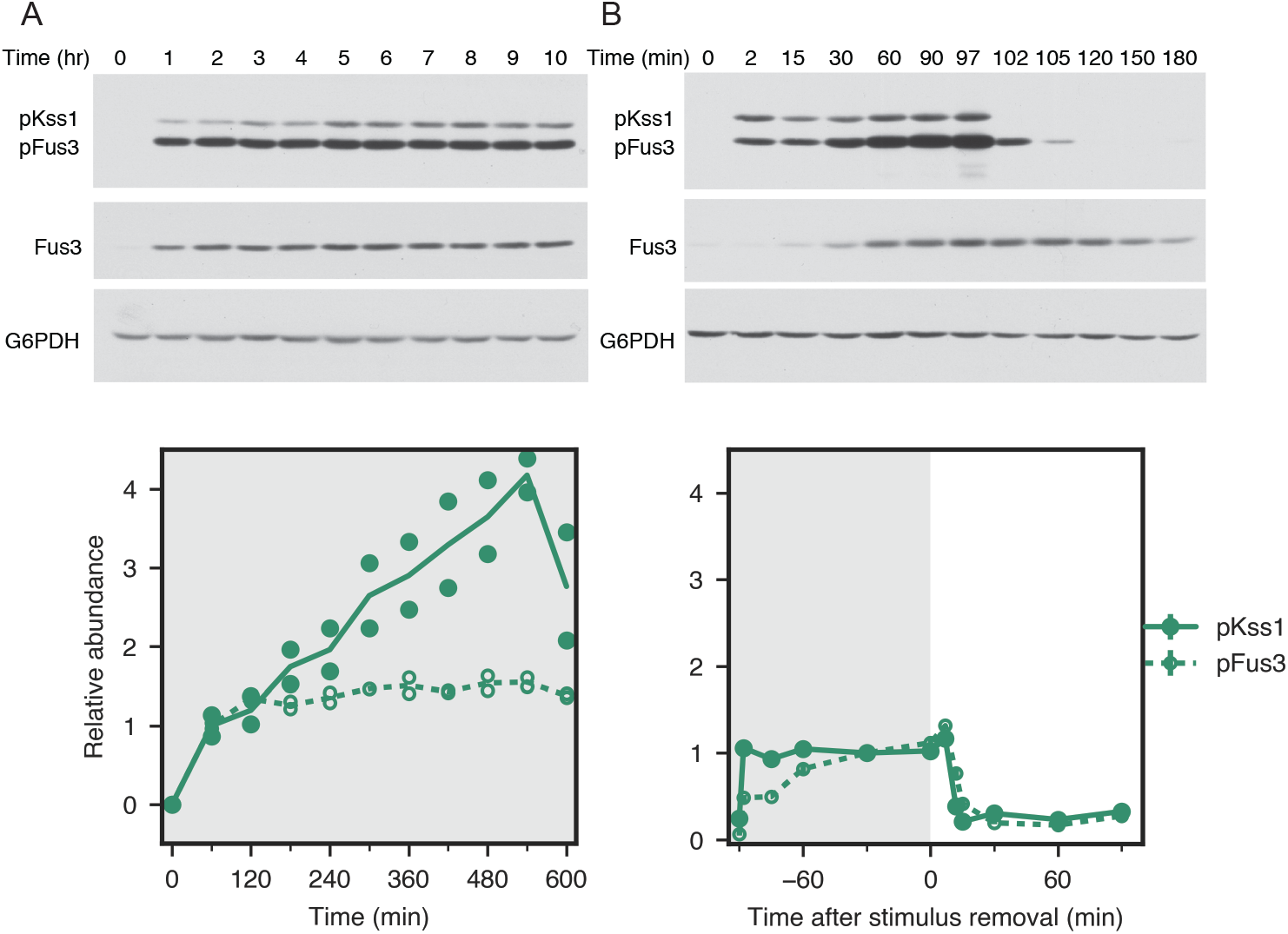
No long-term adaptation or persistence is present in MAPK activation. Quantification of MAPK activation by immunoblotting with phospho-p44/42 MAPK antibodies to detect active MAPK (pKss1 and pFus3), Fus3 antibodies to detect total Fus3, and anti-G6PDH as a loading control for (A) constant stimulus and (B) a 90-minute pulse of stimulus. Quantification of Western blots are presented as either (A) the mean and individual data points from two experiments normalized to the average response after 60 minutes of stimulus exposure or (B) the mean ± standard deviation from three independent experiments. To compare between conditions, quantification of immunoblotting is normalized so pKss1 and pFus3 are equal to 1 after 60 minutes of stimulus.

We similarly monitored Fus3 and Kss1 kinase activity for a 90 min pulse of 50 nM pheromone by Western blot analysis for dual phosphorylation of the MAPKs. In this case aliquots of the culture were removed at indicated intervals during pheromone exposure and after removal of pheromone. Unlike gene expression, activity of the two MAPKs diminished rapidly once pheromone was removed (**Fig. 3B**), demonstrating that the mechanism for persistence also lies downstream of the MAPK signaling molecules.

### Model for transcriptional regulation

To determine which elements of the pathway are critical for regulating the magnitude of the response, long-term adaptation and persistence, we developed a mechanistic model. We chose to include four established signaling motifs. We included an incoherent feedforward loop resulting from Far1-dependent Ste12 degradation as one potential mechanism for long-term adaptation *(22)* (**Fig. 4A, motif 1**). Next, we included positive feedback loops resulting from Ste12 auto-regulation and Ste12-dependent transcription of the MAPKs (**Fig. 4A, motif 2**), which we hypothesized could contribute to both the amplitude and persistence of the signaling response. Previously, we found that if Ste12 in the Dig/Ste12 complex degraded more slowly than free Ste12, rebinding of the Digs to Ste12 could act as a mechanism for adaptation (**Fig. 4A, motif 3**) *(35)*. We hypothesized that this motif also could contribute to persistent signaling, if the rate constant associated with rebinding was small. Finally, we included a negative feedback loop resulting from Ste12 induced synthesis of Far1 (**Fig. 4A, motif 4**), which we hypothesized might also contribute to long-term adaptation. These four motifs were included in the full model to capture persistent activation following stimulus removal and long-term adaptation of the transcriptional response (**Fig. 4B**). Importantly, the model also included synthesis and degradation of our transcriptional reporter. The abundance of this transcriptional reporter was the experimental output used to train the model. The model input was a piecewise-linear function that corresponded to the temporal pheromone stimulation profile. Full details of the mathematical model including the set of differential equations that describe the system are presented in Materials and Methods.

**Figure 4.**
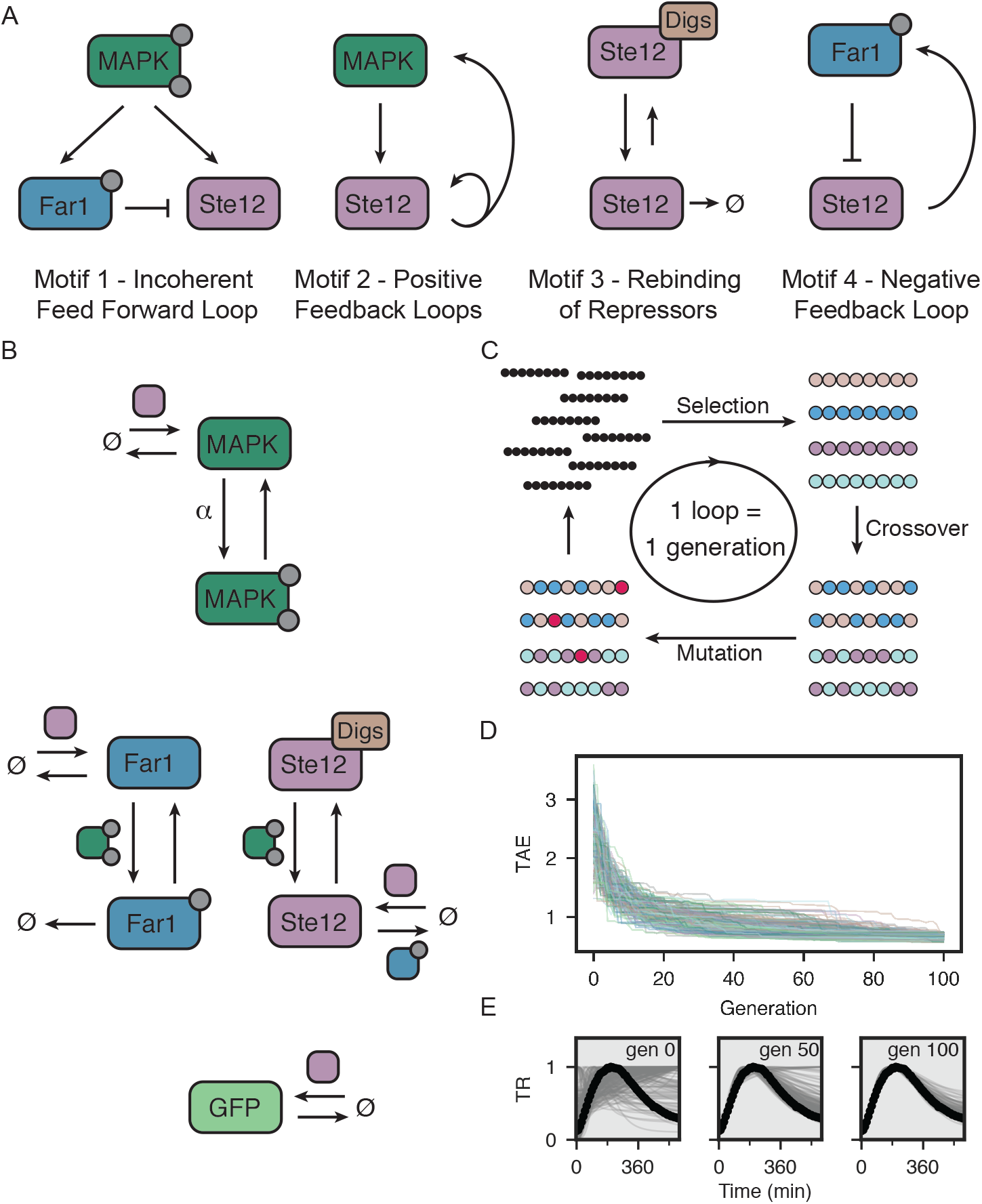
Model of the gene regulatory network. (A) Four important signaling motifs, an incoherent feedforward loop in which phosphorylated Far1 promotes the degradation of Ste12 (Motif 1), positive feedback loops where Ste12 promotes the transcription of itself and the MAPK (Motif 2), slow rebinding of the transcriptional repressors (Dig1 and Dig2) to Ste12 (Motif 3), and a negative feedback where Ste12 promotes the transcription of Far1 (Motif 4). (B) The complete model incorporating all four motifs, which includes MAPK, Far1, and Ste12 each in their active and inactive states as well as the transcriptional reporter GFP. The colored icons adjacent to arrows in the schematic indicate the pathway components that increase each rate. (C) Schematic of evolutionary algorithm (EA) used to fit the model to experimental data. This EA was run 2000 times selecting the best of 500 individuals for 100 generations. (D) The total absolute error (TAE) between the simulation and experimental data for the top 10% of 2000 independent EA runs. Each line represents the lowest error of the 500 individual parameter sets. (E) A comparison of the predicted transcriptional response (TR) of the top 10% of fits at generations 0, 50, and 100 (gray lines) to experimental data (black circles) for constant stimulus.

Below we describe the data sets used to train the model and present results for the model’s performance. We used an evolutionary algorithm (**Fig. 4C**) to fit the model’s 28 parameters. For each parameter, we determined a biologically relevant range from which the parameter values were selected (**Supplementary Material**). Each generation of the evolutionary algorithm had 500 individual parameter sets that underwent selection, crossover, and mutation. Over the course of 100 generations the total absolute error (TAE) between the experimental data and the simulations converged (**Fig. 4D and E**).

### Assessment of model performance

Signaling pathways represent nonlinear dynamical systems capable of responding on multiple different scales. Therefore, we reasoned to develop a predictive model for transcriptional regulation, it was critical to measure the system’s response to time dependent pheromone concentrations with multiple different frequencies. To this end, in addition to the single pulse data described above (replotted in **Figs. 5A-F**), we collected data for periodic stimulation consisting of pulses of pheromone in which the on and off intervals were the same length. The on-off durations used were 45, 60, 75, 90, and 120 (**Fig. 5, G-K**). We also included data for constant pheromone stimulation (replotted in **Fig. 5L**.) The model captured the varying durations of persistence after stimulus is removed (**Fig. 2D**, gray curve) and long-term adaptation to stimulus under conditions of both periodic and constant stimulus (**Fig. 5**). The model was also capable of capturing the dynamics of the MAPK activation profiles in response to constant stimulation and to a 90 min pulse of 50 nM pheromone (**Fig. S2**).

**Figure 5.**
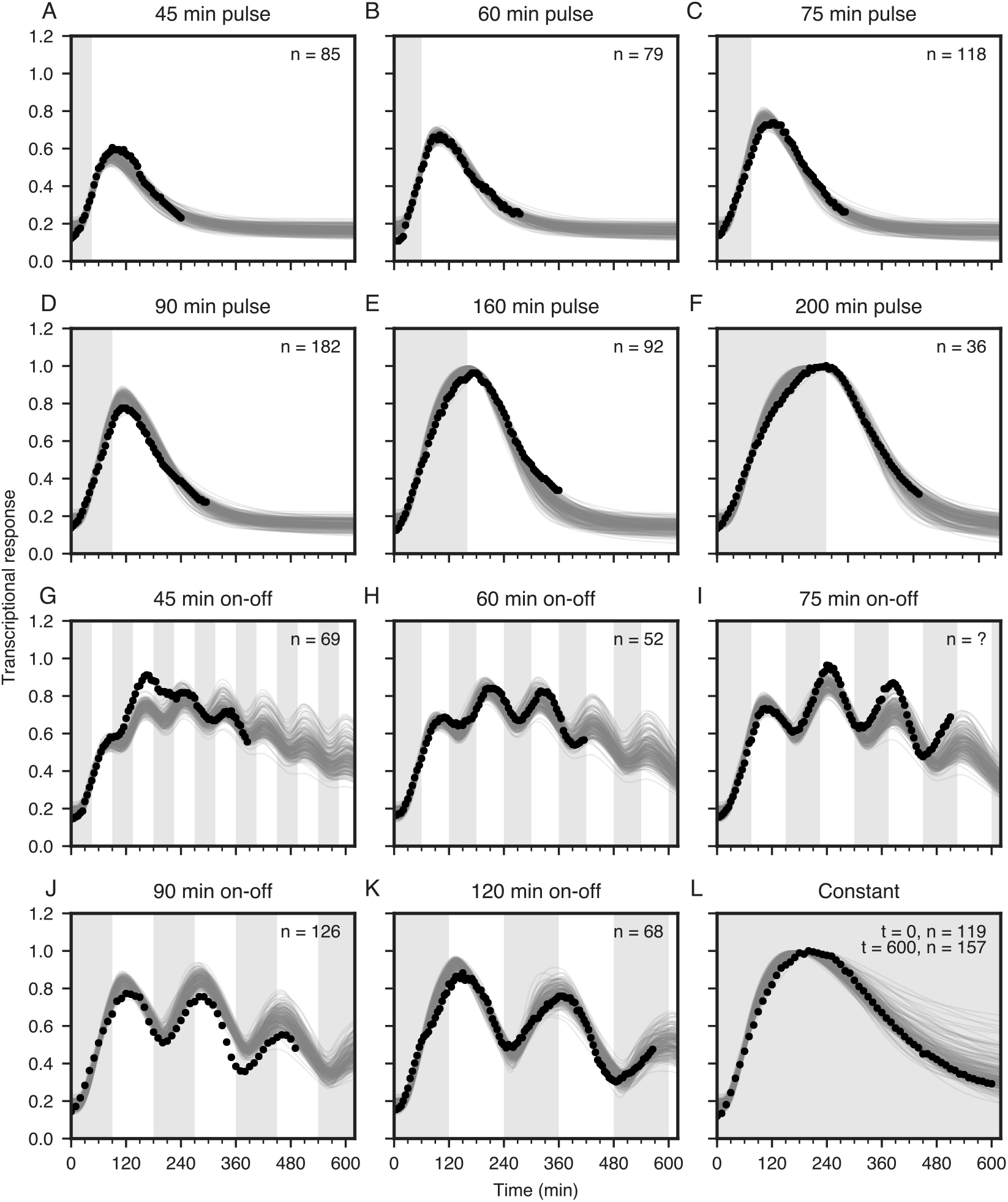
Model captures response to dynamic stimulation. Model simulations generated using the top 10% of parameters found by the evolutionary algorithm (gray lines) compared to the experimental data (circles) for wildtype strain (BY4741-68) transcriptional response to (A-F) six different pulse durations (45, 60, 75, 90, 160, and 200 min), (G-K) five different oscillatory stimulation profiles (45, 60, 75, 90, and 120 min on-off), and (L) constant stimulus of 50 nM pheromone. Gray shading indicates when mating pheromone is present in the time course. Experimental data is presented as the average transcriptional response of the indicated number (n) of cells at the time stimulus is first removed.

### Regulation of response to prolonged stimulus

To understand how response to constant stimulus is regulated, we perturbed motifs that are likely to affect the magnitude of the response and long-term adaptation. First, we used the model to investigate the role of Ste12 auto-regulation in determining the magnitude of the transcriptional response by eliminating motif 2. We found that some parameter sets found by the evolutionary algorithm predict a dampened response in the absence of Ste12 auto-regulation (green and brown curves) while other parameter sets predict a response similar to wildtype (purple curves) (**Fig. 6A**). To experimentally determine whether Ste12 auto-regulation in this system has a significant role in amplifying the response, we replaced the Ste12 promoter with that of the promoter of the scaffold protein Ste5 (*^P^STE5-STE12*). We chose this promoter because it produces constitutive amounts of Ste12 similar to the basal amount from the endogenous promoter (**Fig. S3**) and is not subject to auto-regulation by Ste12 *(36)*. In cells containing the *^P^STE5-STE12* mutation, the overall transcriptional response was diminished, and long-term adaptation began ~50 min sooner than for wildtype cells (**Fig. 6A**, triangles). These findings indicate Ste12 auto-regulation is important for amplifying the response and affects the timing of adaptation. They also suggest that Ste12 autoregulation counterbalances the depletion of Ste12 promoted by Far1 (motif 1).

**Figure 6.**
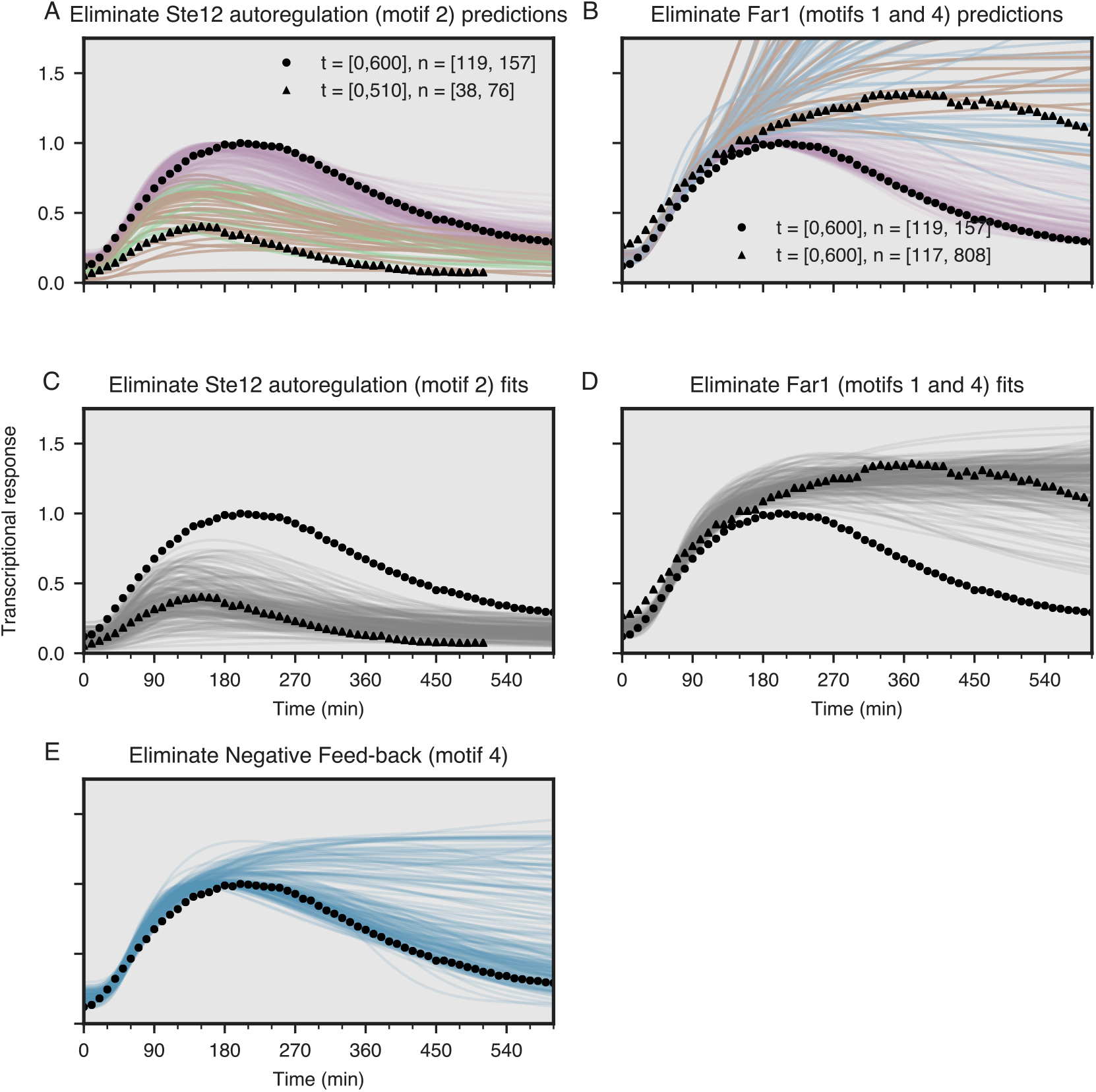
Model predictions of response to sustained stimulus for mutants that perturb signaling motifs. (A and B) Simulations (lines) using the best 10% of parameters found by the evolutionary algorithm for two signaling network perturbations. (A) Simulations in the absence of Ste12 autoregulation (*STE12* endogenous promoter replaced with that from *STE5*) predict a variety of responses ranging from no change (purple lines) from the wildtype (circles) to a dampened response (green and brown lines). Experimental data of the transcriptional response of the *^P^STE5-STE12* mutant (BY4741-103) shows a dampened response (triangles). (B) Simulations in the absence of the incoherent feedforward and negative feedback loops (Far1 removed) predict a variety of responses ranging from no change (purple lines) from the wildtype (circles) to a persistent transcriptional response (brown and blue lines). Experimental data of the transcriptional response of the *far1Δ* mutant (BY4741-130) shows a persistent response (triangles). Parameters from simulations that best capture the experimental results for the *^P^STE5-STE12* mutant (green lines in A) and *far1Δ* mutant (blue lines in B) were selected and used to predict the response of both signaling perturbations (brown lines in A and B). (C and D) Model simulations (gray lines) generated using the top 10% of parameters found by the evolutionary algorithm fit to wildtype (BY4741-68) (constant, single pulse, and periodic stimulus), *far1Δ* (constant stimulus), and *^P^STE5-STE12* (constant stimulus) training data compared to the experimental data for wildtype (circles) or (C) *^P^STE5-STE12* mutant and (D) *far1Δ* mutant responses (triangles). (E) Using the parameter sets shown in C and D, simulations (blue lines) for elimination of only the negative feedback loop. For most parameter sets, elimination of negative feedback exhibits long term adaptation similar to wildtype (circles).

Next, we investigated the role of Far1 in long term adaptation. In the model, Far1 is involved in two signaling motifs. The first is the incoherent feedforward loop (motif 1) formed by MAPK activation of Far1, followed by active Far1 promoting the degradation of Ste12. The second (motif 4) is the negative feedback loop formed by Ste12-dependent expression of Far1. First, we used the model to predict the system’s response when Far1 is eliminated, which blocks both Far1-dependent mechanisms of adaptation (motifs 1 and 4). We found that some parameter sets predict no long-term adaptation in the absence of Far1 (blue and brown curves); however, other parameter sets predict no difference from wildtype (purple curves) (**Fig. 6B**). We reasoned that the interaction between the Digs and Ste12 was responsible for long-term adaptation for those parameter sets still exhibiting adaptation in the absence of Far1 *(35)*. In the model, we allowed for the possibility that complex formation with the Digs protects Ste12 from degradation. There are two consequences of this protective complex *(35)*. First, it ensures a large pool of inactive Ste12 is maintained prior to pheromone stimulation. Second it provides for an adaptive response. The basis for adaptation is that following exposure to pheromone, the free Ste12 concentration transiently increases as Ste12 is released from the Digs, but eventually returns to its pre-stimulus level *(35)*. To test whether eliminating protective binding has any effect on adaptation in our model, we set the degradation rate of Ste12 in the Dig/Ste12 complex equal to that of the degradation rate of free Ste12. With this change, long-term adaptation was lost when the model was run using parameter sets that predicted Far1 was not involved in adaptation (**Fig. S4**). To determine which of these mechanisms is responsible for regulating long-term adaptation, we examined the response of a *far1Δ* mutant strain. In this mutant the transcriptional output does not diminish over time (**Fig. 6B, triangles**) demonstrating that Far1-dependent degradation of Ste12 is the primary mechanism of long-term adaptation.

To further constrain model parameters, we retrained the model including experimental data for the *^P^STE5-STE12* and *far1Δ* mutants. The resulting parameter sets better captured the responses of the pathway mutants than those used for predictions (compare **Figs. 6C and D to Figs. 6A and B**), while maintaining similarly good fits to the wildtype transcriptional responses to different pheromone stimulation regimens (compare **Fig. S5 to Fig. 5**). The distribution of parameters associated with both motifs narrows when the additional data are included in the training sets (compare parameters *kff*, rate of Far1 dependent Ste12 degradation, and *kfb2*, rate of Ste12 autoinduction, in **Fig. S6A and B**). This demonstrates that including strategic pathway perturbations in the training data can improve ability to identify biologically relevant parameters.

Because elimination of Far1 disrupts both the incoherent feedforward and negative feedback motifs (motif 1 and 4, respectively), we used the model to test if negative feedback contributes to long-term adaptation. In the model, disruption of the incoherent feedforward loop is equivalent to eliminating Far1 since promoting degradation of Ste12 is the only effect of Far1 on transcriptional response. However, we can use the model to identify the role of negative feedback. When transcriptional induction of Far1 by Ste12 was eliminated (motif 4) some simulations predict a sustained response, but most simulations still show adaptation (**Fig. 6E**). These results suggest that the incoherent feedforward loop (motif 1) is the predominant mechanism for long-term adaptation of the transcriptional response. While the model did not require the induction of Far1 for long term transcriptional adaptation, it is likely this feedback is required for one of the other functions of Far1 in the mating response, such as gradient sensing or maintaining cell cycle arrest.

### Regulation of response to dynamic stimulus

To examine motifs that could contribute to persistence and further test the model’s predictive power, we measured the system’s response to single 90-minute pulses of 50 nM pheromone in the presence of pathway mutants that perturb Ste12 autoregulation, binding to DNA, or binding to the Dig1 and Dig 2 repressors. First, we eliminated Ste12 autoregulation (motif 2) as before by using the *^P^STE5-STE12* mutation. While there was a dampened response consistent with the response to constant stimulus, neither the simulations nor the mutant response had any appreciable effect on persistence (**Fig. 7A**). Second, we examined a mutation to one of the three pheromone responsive elements (*PREs*) within the *FUS1* promoter that drives transcription of the GFP reporter. Ste12 has been reported to bind at a synthetic promoter having the same *PRE* mutation with only 30% of the affinity that it binds to a synthetic promoter with the wildtype sequence *(37)*. This *PRE* mutation (*PRE*-GFP*) significantly reduced the maximal amplitude of the transcriptional response (**Fig. 7B**, triangles). Using the best 10% of parameter sets found from fitting to the wildtype, *far1Δ*, and *^P^STE5-STE12* data, we predicted the response of the *PRE* mutant by increasing the apparent dissociation constant by a factor of 3.33. The resulting parameter sets accurately predict the response of the *PRE*-GFP* mutant (**Fig. 7B**).

**Figure 7.**
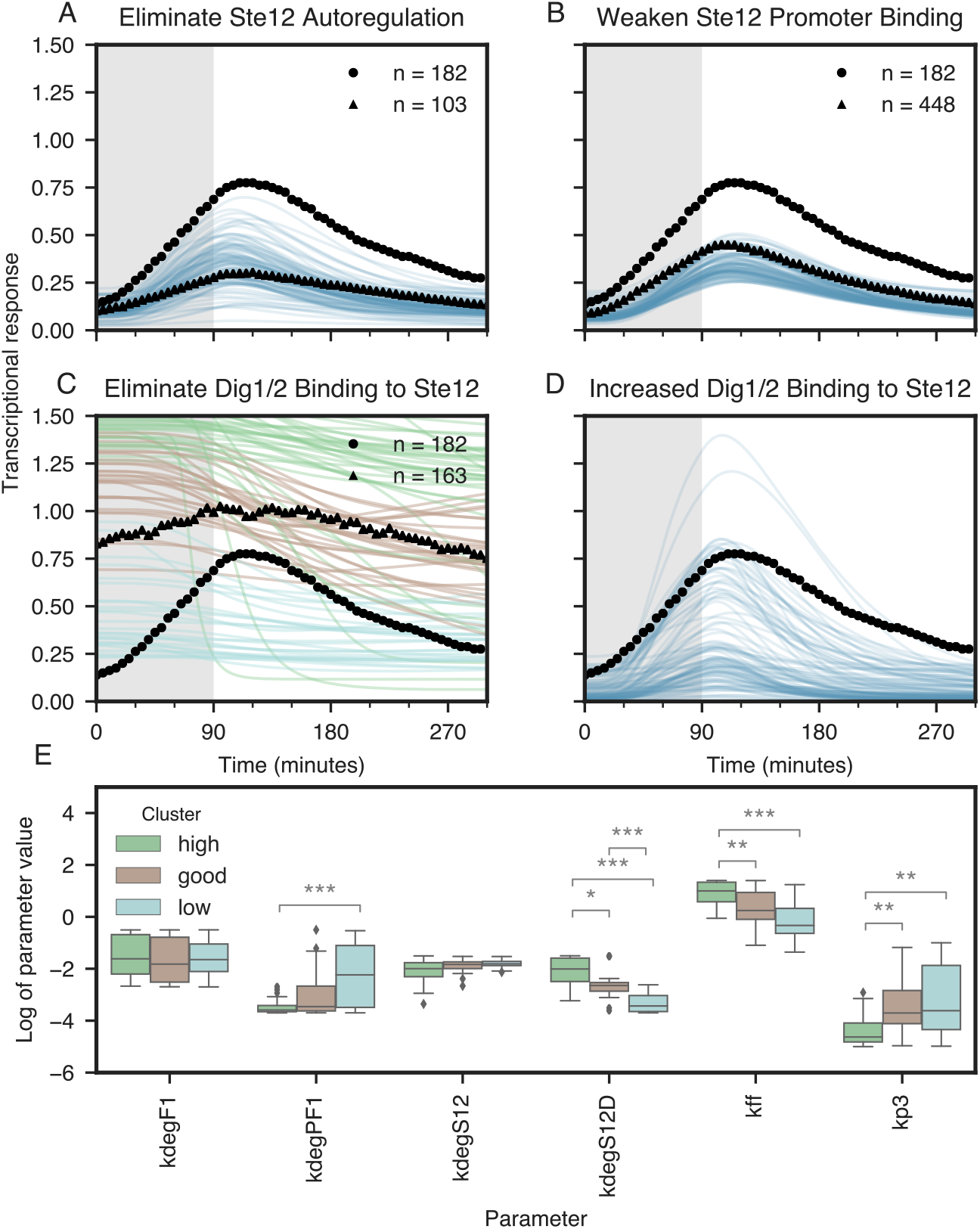
Prediction response of pathway perturbations to transient stimulus. The top 10% of parameter sets found by the evolutionary algorithm fit to wildtype (BY4741-68), *far1Δ* (BY4741-130), and *^P^STE5-STE12* (BY4741-103) training data were used to predict the response of pathway perturbations to a 90-minute pulse of stimulus. (A) Predicted response of a the *^P^STE5-STE12* mutation that eliminates autoregulation of Ste12 (blue lines) compared to experimentally determined response for the mutant strain (triangles). (B) Predicted response of a *PRE*-GFP* promoter mutation that causes Ste12 to bind less tightly to the GFP promoter (blue lines) compared to experimentally determined response for the mutant strain (BY4747-169) (triangles). (C) Predicted response of a *dig1Δdig2Δ* double mutation that eliminates the transcriptional repressors divided into three clusters, high basal response (green lines), low basal response (cyan lines), and response that best fits the experimental data (brown lines) compared to experimentally determined response for the mutant strain (BY4741-147) (triangles). (D) Predicted response of faster rebinding of the transcriptional repressors (Dig1 and Dig2) (blue lines) compared to wildtype response (circles). Wildtype response (circles) is included on all panels A-D as a reference. (E) Analysis of parameter distributions within each of the clusters shown in panel C for the rates of inactive Far1 degradation (kdegF1), active Far1 degradation (kdegPF1), active Ste12 (kdegS12), Ste12 in complex with the transcriptional repressors (kdegS12D), Far1 dependent degradation of Ste12 (kff), and dephosphorylation of active Far1 (kp3). Significance values (*p < 0.5, **p < 0.1, and ***p <0.01) were calculated using a t-test with a Bonferroni correction for multiple hypothesis testing. Similar analysis for all parameters is available in the supplement (Fig. S7).

Finally, we examined the effect deleting the Dig1 and Dig2 repressors, which causes Ste12 to be constitutively active. This deletion mutant showed high basal expression and a slight increase in expression following pheromone induction (**Fig. 7C, triangles**). We predicted the response of the *dig1Δdig2Δ* mutant by setting the total Dig concentration to zero. The model predicted a wide range of responses in the absence of the Dig1 and Dig2 transcriptional repressors. Interestingly, the results could be clustered into three groups (**Fig. 7C, colored curves**). The model predictions that showed high basal transcriptional response (**Fig. 7C, green curves**) result from parameter sets in which the degradation of Ste12 in complex with the Digs (*kdegS12D*) is similar to that of the degradation rate of free Ste12 (*kdegS12*) (**Fig. 7E**). In this case the total amount of Ste12 is the same in the *dig1Δdig2Δ* mutant and wildtype reference. Removing the Dig repressors generates more active Ste12 prior to pheromone stimulation, and, therefore, higher levels of the reporter in the mutant. For model predictions in which the prestimulation level of the fluorescent reporter does not increase significantly compared to the wildtype reference (**Fig. 7C, cyan curves**), removing the Digs had two effects. For these parameter sets, the degradation rate of Ste12 in complex with the Digs is reduced (**Fig. 7E**). That is, the Dig repressors provide protective binding. Removing the Digs exposes Ste12 for degradation, but also activates Ste12. When these two opposing effects are balanced, the pre-stimulation level of active Ste12 in the *dig1Δdig2Δ* mutant is similar to that of wildtype, and, therefore, the expression level of the reporter does not significantly increase. The parameter sets that fit the experimental data best had intermediate degradation rates for Ste12 in complex with the Digs (**Fig. 7E** and **Fig. 7C, brown curves**). These results are consistent with our previous analysis of Ste12 dynamics that demonstrated that the Dig repressors provide some degree of protective binding *(35)*.

Another observation consistent with previous experimental observations is that parameter sets best fitting the experimental results for the *dig1Δdig2Δ* mutant (**Fig. 7C**, **brown curves**) predict that the rate at which active Far1 is degraded (*kdegPF1*) is less than that for inactive Far1 (*kdegF1*) (**Fig. 7E**) *(38)*. Additionally, other parameters that affect Far1-dependent degradation of Ste12 including the rate of Far1 dephosphorylation (*kp3*) and rate of Far1 dependent degradation of Ste12 (*kff*) show significantly different ranges for the three groups of parameter sets (**Fig. 7E**). The best fitting predictions show modest attenuation of the transcriptional response resulting from the feedforward Far1-dependent Ste12 degradation, consistent with the experimental results (**Fig. 7C**, **brown curves**). The high responders (**Fig 7C, green curves**) have parameter values that increase the abundance of Far1 resulting in a stronger effect of the incoherent feedforward and consequently predictions of stronger transcriptional attenuation. Conversely, the low responders (**Fig. 7C, cyan curves**) have parameter values that rapidly degrade and deactivate active Far1 both of which reduce the effect of the incoherent feedforward and consequently predict little to no transcriptional attenuation. These results again illustrate the need to use targeted pathway perturbations to fully constrain model parameters.

Because in the model, the only mechanism for transcriptional induction is dissociation of Ste12 from the Dig repressors, the model is not able to capture the slight pheromone-dependent induction seen in the *dig1Δdig2Δ* strain. This induction may result from pheromone-induced degradation of the transcription factor Tec1, a known binding partner of Ste12 *(39)*. The slight pheromone-dependent induction in the *dig1Δdig2Δ* strain exhibits prolonged maximal expression after a 90-minute pulse of stimulus (72 min persistence) compared to wildtype (43 min persistence). To further investigate how the transcriptional repressors contribute to persistence, we perturbed motif 3 by increasing the rebinding rate of Ste12 to the Digs in the model by 5-fold. In doing so, the average persistence of the simulations decreased from 25 min to 13 min (**Fig. 7D**). This result combined with the prolonged persistence when the transcriptional repressors are deleted suggests that slow rebinding of the transcriptional repressors are a primary factor in the persistent transcriptional response following stimulus removal.

### Prediction of different stimulation profiles

To further test the model’s predictive power, we measured the response of cells exposed to periodic stimulation at the same frequencies as shown in Fig. 5, but at a lower pheromone concentration (**Fig. 8**, triangles). In response to 10 nM of constant pheromone, the fluorescent reporter achieves the same maximum amplitude as the 50 nM case but takes 25 min longer to reach its half maximum amplitude (**Fig. 8A**). For short pulses of stimulus (**Fig. 8B**), the amplitude of the response to 10 nM is considerably lower than that to 50 nM for all pulses. However, for longer pulses (**Fig. 8C**) there is less of a difference in the amplitude between the two doses. To simulate the lower pheromone dose, the only modification we made to the model was to adjust the slope of the input signal to match the slower production rate of the fluorescent reporter measured at 10 nM constant pheromone. Using this adjustment to the input stimulus, all of the parameter sets that fit the 50 nM data accurately predicted the response to sustained and pulsed 10 nM pheromone (**Fig. 8**, blue curves). This performance demonstrates that this model is capable of capturing behaviors at different doses of stimulus despite only being trained on a single dose.

**Figure 8.**
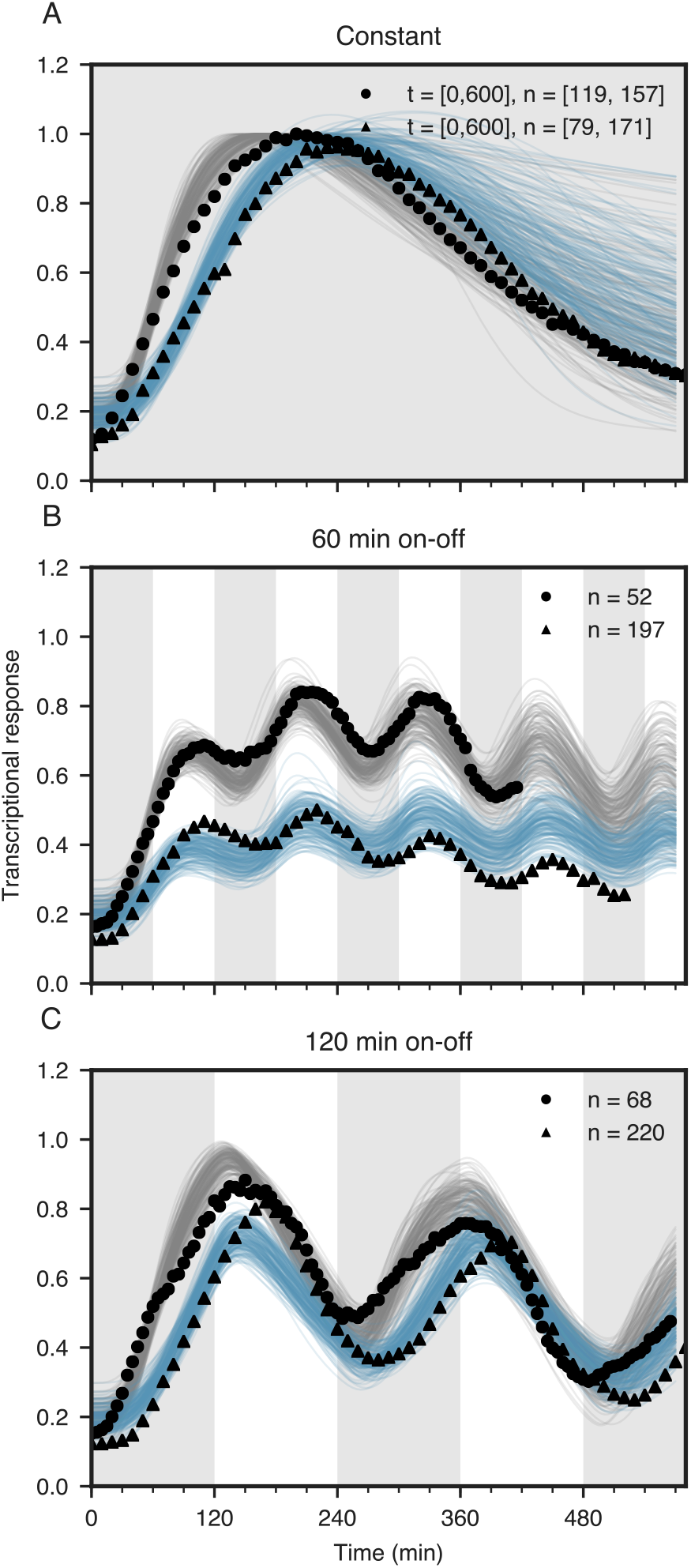
Prediction of response to a different dose of stimulus. The top 10% of parameter sets found by the evolutionary algorithm fit to all training data from wildtype (BY4741-68), *far1Δ* (BY4741-130), and *^P^STE5-STE12* (BY4741-103) strain responses to 50 nM pheromone (gray lines) were used to predict the response of the wildtype strain (BY4741-68) to 10 nM pheromone (blue lines) for (A) constant and (B and C) two periodic stimulation profiles (60 and 120 min on-off). Experimental data for 50 nM stimulus is represented by circles and experimental data for 10 nM stimulus is represented by triangles.

## DISCUSSION

A common way for cells to respond to changes in their environment is by regulating gene expression. Because environmental conditions are dynamic and can show significant variability, gene expression needs to be tightly regulated by the signaling pathways used by cells to monitor their surroundings. This regulation, which typically takes the form of feedback and feedforward loops, makes gene regulation an inherently non-linear process. Therefore, predicting the response of these systems is not possible without the aid of mathematical models. Developing predictive models is challenging for two reasons: 1) these models tend to contain many parameters that are not directly measurable and therefore must be estimated from experimental data and 2) these systems operate on multiple time scales and, therefore, experimental data sets used to train the models must capture the relevant time scales. To overcome both these obstacles, we developed a research strategy that involved exposing yeast cells to single and periodic pulses of mating pheromone. By varying both the duration and frequency of the pulses, we ensured that the regulatory network that controls gene expression during the mating response was probed on the relevant time scale and with sufficient temporal resolution to accurately perform parameter estimation. This systematic analysis revealed novel features of the pathway and allow us to develop a mathematical model with predictive power.

Our analysis led us to the discovery of memory in the yeast mating response. Specifically, we discovered that transcriptional regulation was sustained following removal of pheromone, a property of the system that we refer to as “persistence”. Our model revealed that this persistence was not due to positive autoregulation of Ste12 but rather involves slow rebinding of the transcriptional repressors to Ste12. Persistent signaling may represent an important design feature of the pheromone response pathway. Yeast mating takes place in noisy environments where pheromone levels are expected to fluctuate. Preparing for mating takes a significant fraction of the cell’s resources. Therefore, once the decision has been made to commit to the mating, it is important that the cell not “give up” if there is a transient loss of the pheromone signal. Persistent signaling provides a mechanism to guard against this situation. Conversely, it is also important that a cell not remain committed to mating indefinitely. This might explain why persistent gene expression does not rely on positive feedback, which is capable of generating an irreversible switch.

In the presence of sustained pheromone signals, it is probably beneficial for yeast cells not to remain growth arrested when mating is unlikely to be successful. Such adaptative behavior in the mating response has been observed previously *(22, 40)*. However, these studies were done in the presence of the protease Bar1, which degrades pheromone, thus making it difficult to identify the predominate mechanism that underlies transient signaling. Interestingly, our results revealed that in the absence of Bar1, pheromone-induced gene expression is transient, whereas MAPK signaling is sustained. Our model predicted two mechanisms could underlie this long-term adaptation, an incoherent feedforward loop or protective binding. Further experiments revealed that the incoherent feedforward loop involving Far1-dependent degradation of Ste12 accounts for most of the long-term adaptation.

Our approach that combines mathematical modeling with experiments designed to probe cellular response pathways over multiple time scales provides a general framework for investigating gene regulatory motifs. First, we used experiments to narrow the portion of the pathway responsible for the dynamic properties of interest. In our case, these preliminary investigations revealed that both persistent signaling and long-term adaptation occurred at the level of gene regulation and did not involve upstream signaling components. Next, we developed a model incorporating known regulatory mechanisms and narrowed parameter ranges to physiologically relevant values. We then performed parameter estimation using an evolutionary algorithm applied to training data sets spanning multiple timescales. The use of time-dependent stimuli covering multiple time scales was essential for building a predictive model. When a subset of the data was used, model parameters were significantly less constrained, and the model’s predictive power was reduced. Additionally, training on data sets spanning multiple timescales revealed the differences in timing of the signaling motifs. For example, the rebinding of the transcriptional repressors and the incoherent feedforward operate on different timescales, leading to decreased persistence after longer pulses of stimulus.

We also note that successful model building is an iterative process. For example, when fit only using wildtype data the model found two mechanisms of long-term adaptation were consistent with the data. The model also predicted positive feedback contributed to amplifying the signal but showed significant variability in the predicted strength of this feedback. Strategic experiments using targeted mutants were then able to identify the true mechanism of long-term transcriptional adaptation and quantify the role of positive feedback. Including these results in the training data sets, further constrained parameter values and allowed the model to accurately predict the system’s behavior for lower pheromone concentrations and additional genetic perturbations.

Because gene editing and quantitative experimental approaches are becoming increasingly more feasible in other cell types, including mammalian cells, we believe our approach can be adapted to these systems. For example, such studies could reveal important information about the dynamics of MAPK signaling pathways dysregulated in diseases, including cancer, and ultimately suggest treatments for restoring proper function.

## MATERIALS AND METHODS

### Plasmids, PCR alleles, and recombinant DNA procedures

**Table 1** lists plasmids used in this study. Those that have been described previously are listed with the corresponding reference. Standard recombinant DNA procedures were used for construction of those plasmids described below *(41)*. **Table 2** lists the sequence of oligonucleotides used for PCR fragment amplification, mutagenesis, and DNA sequence confirmation involved in plasmid and strain constructions.

**Table 1.**
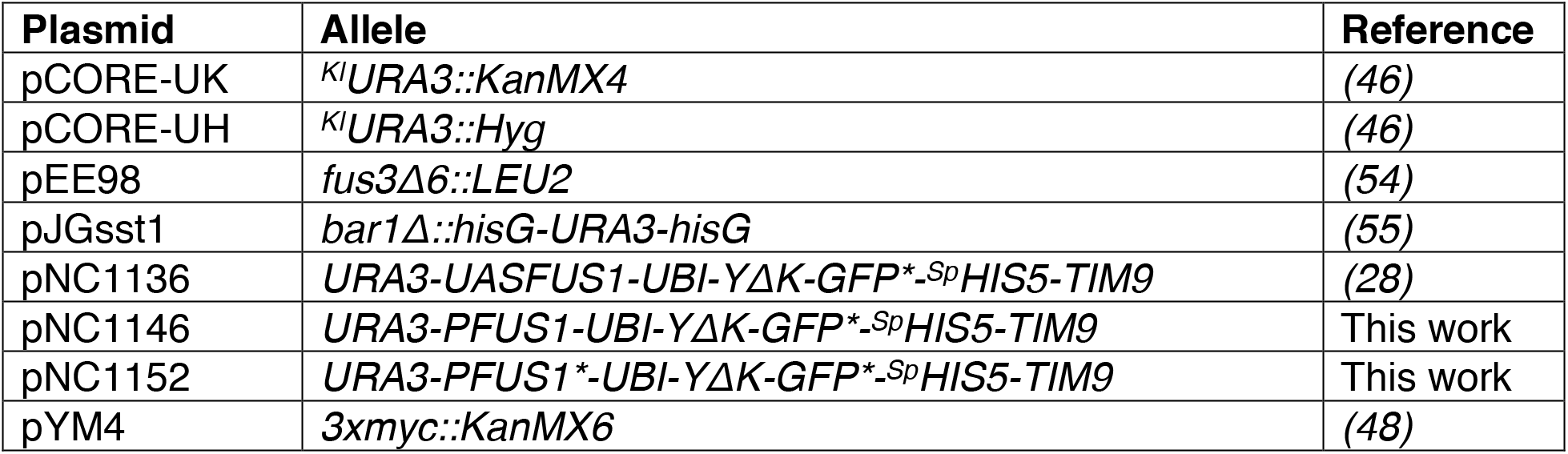
Plasmids.

| **Plasmid** | **Allele** | **Reference** |
| --- | --- | --- |
| pCORE-UK | *KlURA3::KanMX4* | (46) |
| pCORE-UH | *KlURA3::Hyg* | (46) |
| pEE98 | *fus3Δ6::LEU2* | (54) |
| pJGsst1 | *bar1Δ::hisG-URA3-hisG* | (55) |
| pNC1136 | *URA3-UASFUS1-UBI-YΔK-GFP\*<sup>-Sp</sup>HIS5-TIM9* | (28) |
| pNC1146 | *URA3-PFUS1-UBI-YΔK-GFP\*<sup>-Sp</sup>HIS5-TIM9* | This work |
| pNC1152 | *URA3-PFUS1\*-UBI-YΔK-GFP\*<sup>-Sp</sup>HIS5-TIM9* | This work |
| pYM4 | *3xmyc::KanMX6* | (48) |

**Table 2.**
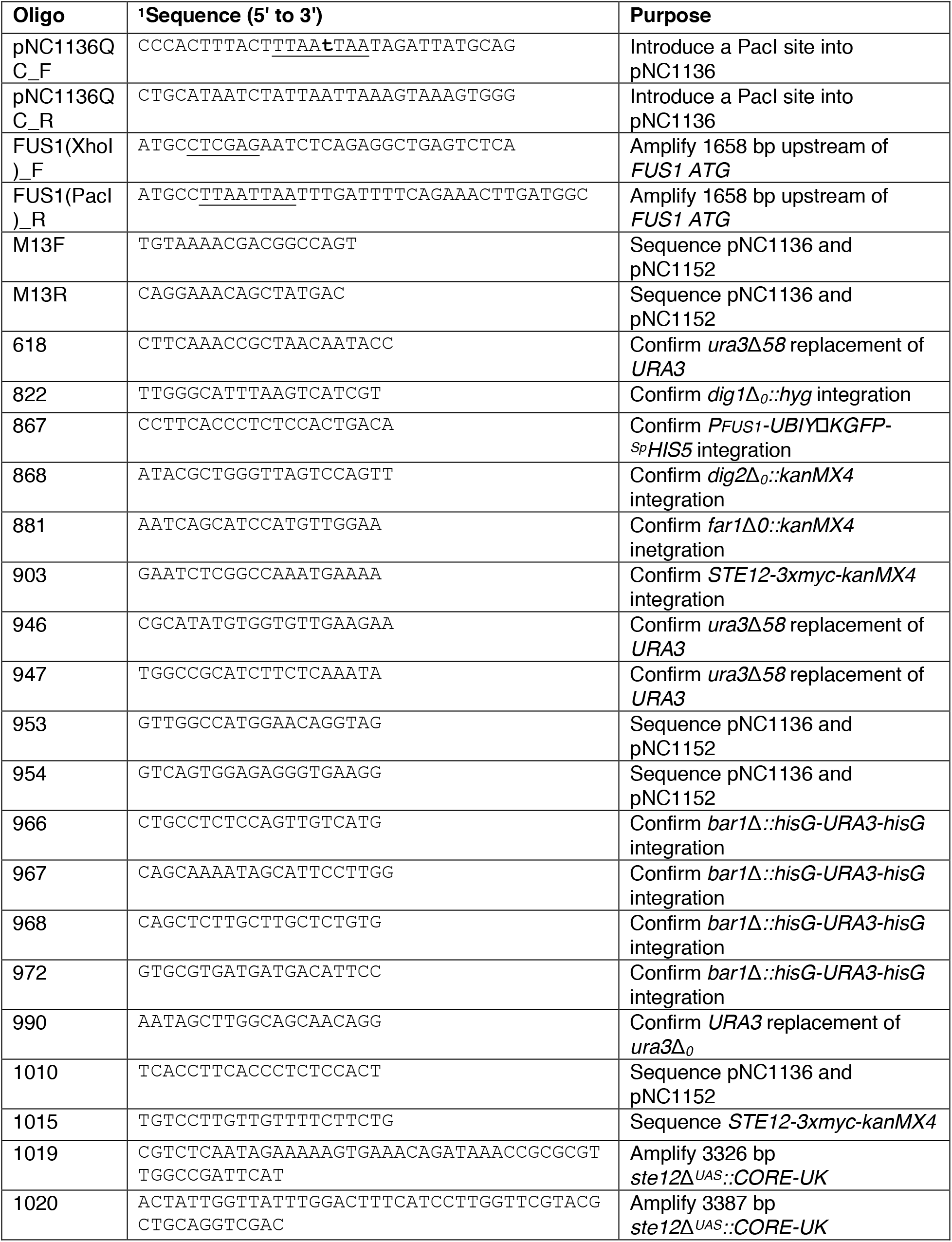
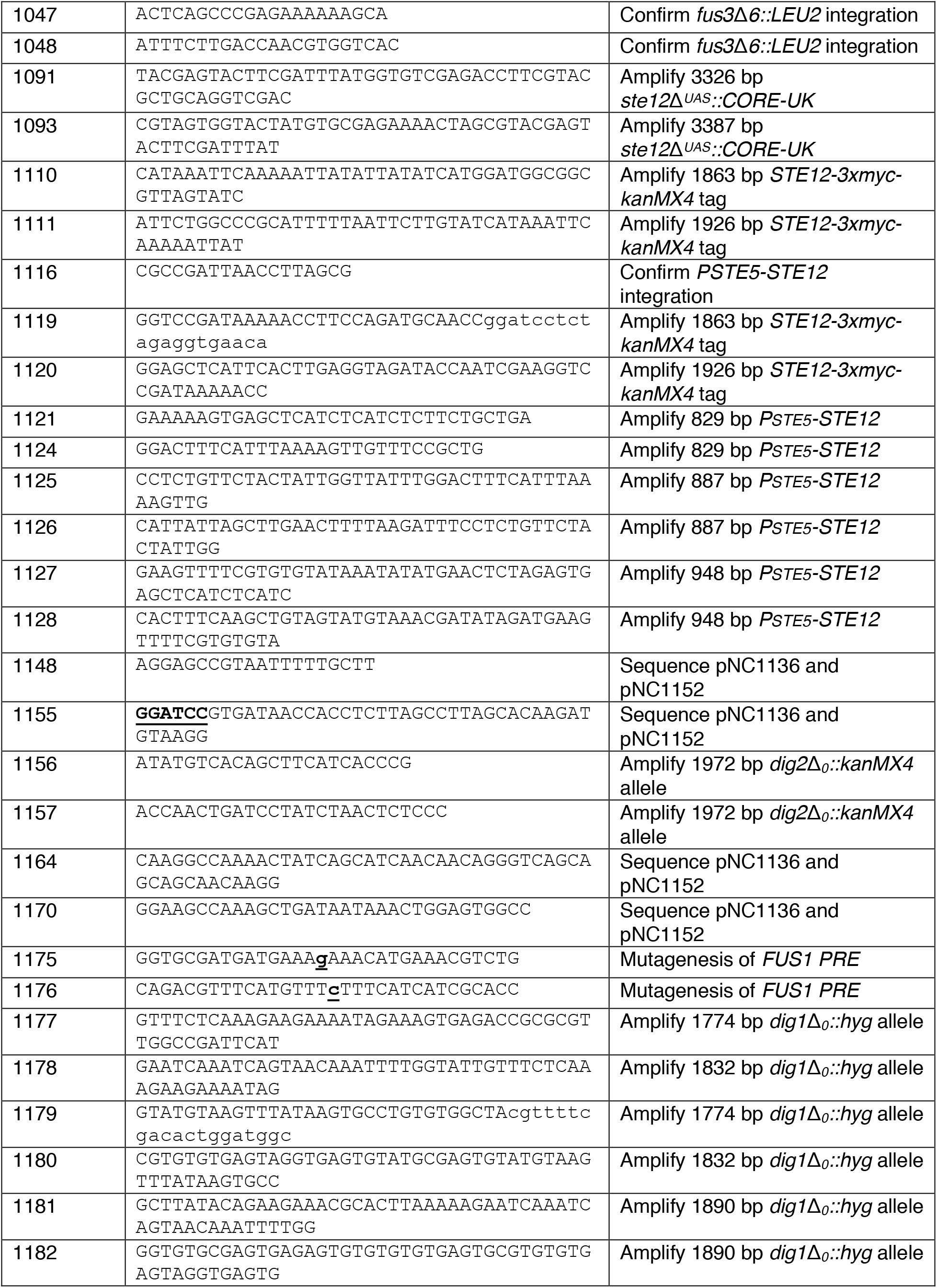
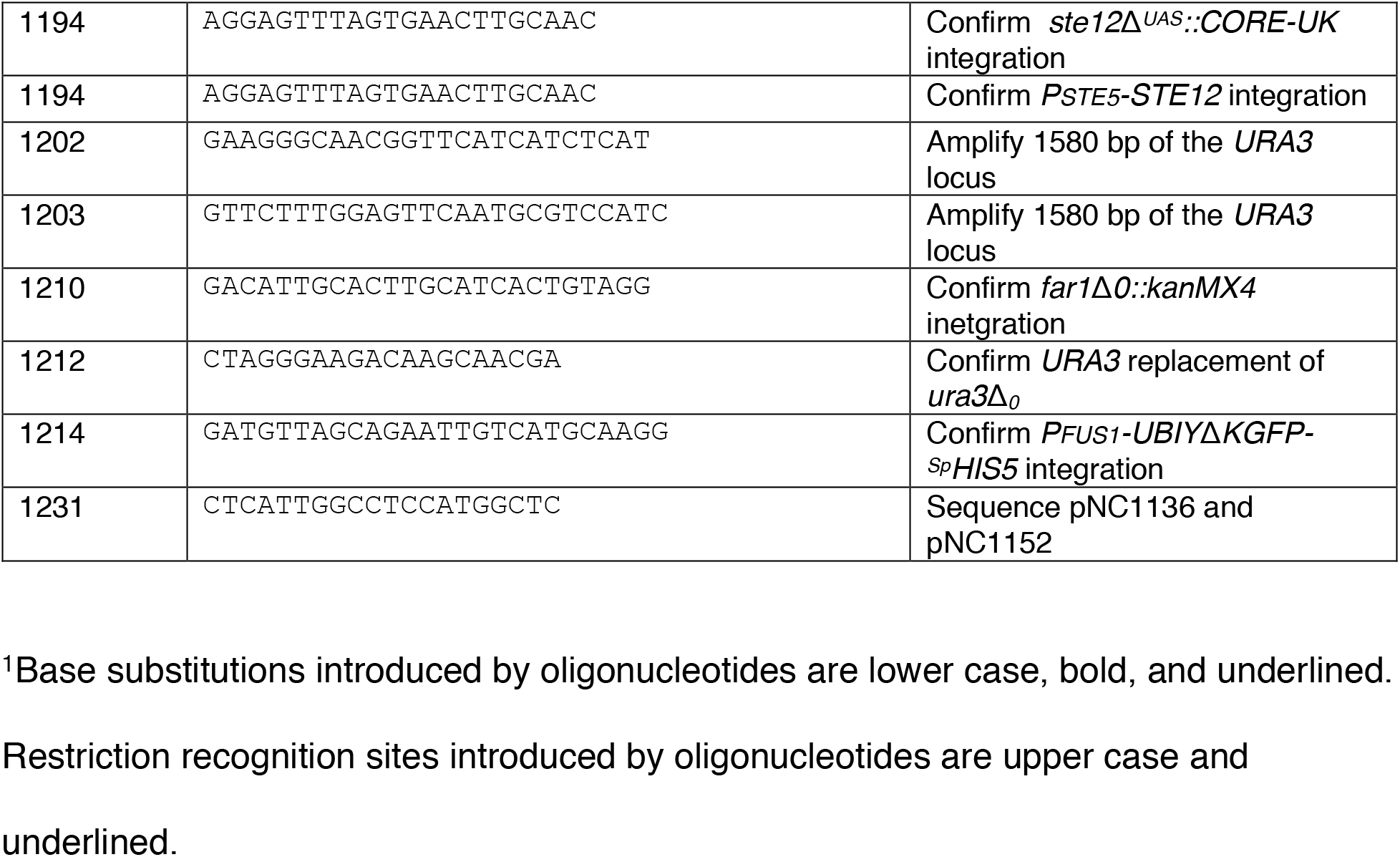
Oligonucleotides.

A fluorescent protein (GFP*) with a fast maturation time and an N-degron tag (Ubi-YΔK) that confers a short half-life was designed and characterized previously *(28)*. The plasmid pNC1146 carries a reporter gene in which the pheromone responsive *FUS1* promoter (*^P^FUS1*) drives expression of the *UBIY*Δ*KGFP\** reporter gene in a cassette with the *S. pombe* (*Sp*) *HIS5* gene as a selectable marker and flanking sequences that target integration to the *URA3-TIM9* intergenic region. To construct this reporter cassette (*URA3-^P^FUS1-UBIY*Δ*KGFP*-^Sp^HIS5-TIM9*), we first introduced a PacI restriction endonuclease recognition site 6 bp upstream of the ubiquitin (*UBI*) coding sequence in the plasmid pNC1136 *(28)*. This modification was accomplished using the Stratagene Quick Change site-directed mutagenesis protocol (Stratagene, La Jolla, CA) with pNC1136 as template DNA and oligonucleotides pNC1136QC(PacI)_F and pNC1136QC_R as primers. Next, a 1658 bp fragment encompassing the *FUS1* promoter flanked by XhoI and PacI restriction endonuclease recognition sites was PCR amplified using BY4741 genomic DNA as template and oligonucleotides FUS1(XhoI)_F and FUS1(PacI)_R as primers. pNC1136 modified with the PacI site and the PCR amplified DNA fragment were digested with XhoI and PacI. The resulting 1646 bp XhoI-PacI *FUS1* promoter fragment (*^P^FUS1*) was ligated to the 6553 bp XhoI-PacI fragment from the plasmid to generate pNC1146. DNA sequence analysis of pNC1146 using primers M13R, 1155, 1164, 1170, and 1231 confirmed the absence mutations in the *FUS1* promoter region.

The plasmid pNC1152 (*URA3-^P^FUS1*(*PRE\**)-*UBIY*Δ*KGFP*-^Sp^HIS5-TIM9*) has the same reporter gene cassette as described for pNC1146 except for a single base pair substitution (C:G to g:c) in one of the *PRE* elements (underlined) that comprise the *^P^FUS1* upstream activating sequence (UAS): <u>ATGAAACA</u>AAC<u>ATGAAACGT</u>CTGTAAT<u>TTGAAACA</u> to <u>ATGAAAgA</u>AAC<u>ATGAAACGT</u>CTGTAAT<u>TTGAAACA</u>. This transversion substitution in the consensus *PRE* was shown by Su et al. *(19)* to shift the equilibrium towards less favorable binding to Ste12. The substitution mutation in the reporter gene cassette was generated using the Stratagene Quick Change protocol (Stratagene, La Jolla, CA) for site-directed mutagenesis with pNC1146 DNA as template, oligonucleotides 1175 and 1176 as primers and Phusion High Fidelity Polymerase (Thermo Scientific, Pittsburgh, PA). DNA sequence analysis of the 5141 bp region encompassing the reporter gene cassette in the isolate designated pNC1152 using oligonucleotide primers M13R, M13F 491, 953, 954, 1010, 1148, 1155, 1170, and 1231 confirmed the presence of the desired mutation in the *P_FUS1_ UAS* and the absence of any additional mutations.

### Yeast strains and genetic procedures

**Table 3** lists yeast strains used in these studies. Media preparation and standard yeast genetic methods for transformation, gene replacement, crosses and tetrad dissection were as described in Amberg, Burke, and Strathern *(42)*.

**Table 3.** Strains used in this study.

| Strain | Genotype | Reference or Source<br>Purpose (Model Perturbation) |
| --- | --- | --- |
| BY4741 | *MATa his3Δ1 leu2Δ0 met15Δ0 ura3Δ0* | Parent strain (56) |
| BY4741-29 | *MATa his3Δ1 leu2Δ0 met15Δ0 ura3Δ0 dig2Δ<sub>0</sub>::kanMX4* | Yeast knockout collection (Invitrogen, Carlsbad, CA)<br>Source of *dig2Δ::kanMX6* allele |
| BY4741-64 | *MATa his3Δ1 leu2Δ0 met15Δ0 URA3* | This work<br>Restores *URA3* targeting region in the BY4741 background |
| BY4741-65 | *MATa his3Δ1 leu2Δ0 met15Δ0 ura3Δ58* | This work<br>58 bp *Apal*-*Stul* deletion in the *URA3* coding sequence |
| BY4741-66 | *MATa his3Δ1 leu2Δ0 met15Δ0 ura3Δ58 bar1Δ::hisG* | This work<br>Eliminates Bar1 protease |
| BY4741-68 | *MATa his3Δ1 leu2Δ0 met15Δ0 ura3Δ58 bar1Δ::hisG PFUS1-UBIYΔKGFP-SpHIS5* | This work<br>Wild-type reference *PFUS1-UBIYΔKGFP* reporter strain; Precursor to BY4741-93, -103, -105, -110, -130, -132, and 147 |
| BY4741-70 | *MATa his3Δ1 leu2Δ0 met15Δ0 ura3Δ0 far1Δ<sub>0</sub>::kanMX4* | Yeast knock out collection (Invitrogen, Carlsbad, CA)<br>Source of *far1Δ<sub>0</sub>::kanMX4* allele |
| BY4741-100 | *MATa his3Δ1 leu2Δ0 met15Δ0 ura3Δ58 bar1Δ::hisG PSTE12::CORE-UK::STE12 PFUS1-UBIYΔKGFP-SpHIS5* | This work<br>Precursor to BY4741-103 |
| BY4741-103 | *MATa his3Δ1 leu2Δ0 met15Δ0 ura3Δ58 bar1Δ::hisG PSTE5-STE12 PFUS1-UBIYΔKGFP-SpHIS5* | This work<br>Eliminates positive feedback (Fig. 4, motif 2) |
| BY4741-105 | *MATa his3Δ1 leu2Δ0 met15Δ0 ura3Δ58 bar1Δ::hisG STE12-3xmyc::KanMX4 PFUS1-UBIYΔKGFP-SpHIS5* | This work<br>Western blot analysis to establish basal *STE12-3xmyc* expression |
| BY4741-110 | *MATa his3Δ1 leu2Δ0 met15Δ0 ura3Δ58 bar1Δ::hisG dig2Δ::kanMX4 PFUS1-UBIYΔKGFP::SpHIS5* | This work<br>Precursor to BY4741-147 |
| BY4741-112 | *MATa his3Δ1 leu2Δ0 met15Δ0 ura3Δ0 bar1Δ::hisG-URA3-hisG* | This work<br>Precursor to BY4741-114 |
| BY4741-114 | *MATa his3Δ1 leu2Δ0 met15Δ0 ura3Δ0 bar1Δ::hisG* | This work<br>Precursor to BY4741-120 |
| BY4741-120 | *MATa his3Δ1 leu2Δ0 met15Δ0 URA3 bar1Δ::hisG* | This work<br>Precursor to BY4741-122 |
| BY4741-122 | *MATa his3Δ1 leu2Δ0 met15Δ0 ura3Δ58 bar1Δ::hisG* | This work<br>Precursor to BY4741-137, -148 and -152 |
| BY4741-130 | *MATa his3Δ1 leu2Δ0 met15Δ0 ura3Δ58 bar1Δ::hisG*<br>*PFUS1-UBIYΔKGFP-SpHIS5 far1Δ::kanMX4* | This work<br>Eliminates Fus3 and Far1 dependent negative feed forward (Fig. 4, motif 1) |
| BY4741-132 | *MATa his3Δ1 leu2Δ0 met15Δ0 ura3Δ58 bar1Δ::hisG*<br>*PFUS1-UBIYΔKGFP-SpHIS5 PSTE5-STE12-3xmyc::KanMX4* | This work<br>Western blot analysis to establish basal *PSTE5-Ste12-3xmyc* expression |
| BY4741-137 | *MATa his3Δ1 leu2Δ0 met15Δ0 ura3Δ58 bar1Δ::hisG*<br>*PFUS1-UBIYΔKGFP-SpHIS5* | This work<br>Wild-type reference reporter strain |
| BY4741-147 | *MATa his3Δ1 leu2Δ0 met15Δ0 ura3Δ58 bar1Δ::hisG*<br>*dig1Δ::hyg dig2Δ::kanMX4 PFUS1-UBIYΔKGFP::SpHIS5* | This work<br>Eliminate repressor inactivation of Ste12 (Fig. 4, Motif 3) |
| D502-3C | *MATΔ ade6* | F. Sherman (University of Rochester, Rochester, NY) |

#### Strains constructed using the one step gene replacement method (43)

<u>*URA3* strain BY4741-64</u> was derived from *ura3*Δ_*0*_ strain BY4741 by transformation with a 1580 bp fragment that was PCR amplified using primer pair 1202/1203 with genomic DNA from the *URA3* strain D502-3C as template and selection on -Ura medium.

<u>*ura3*Δ*58* strain BY4741-65</u> was derived from BY4741-64 by replacing the *URA3* allele with the HindIII fragment from pURA3Δ58 (provided to us by M. Resnick, NIEHS) and selecting for the resulting Ura-phenotype using 5-FOA medium. This *ura3*Δ*58* null allele has a 58 bp deletion of an ApaI-StuI fragment in the *URA3* coding sequence. PCR analysis confirmed the 58 bp deletion by using BY4741-65 genomic DNA as template with primer pair 946/618, which fail to yield a product, and primer pair 946/947, which yield a smaller product than the for the wildtype reference strain.

<u>*bar1*Δ::*hisG* strainBY4741-66</u> was derived from BY4741-65, by using the EcoRI -SalI fragment from pJGsst1 to replace the *BAR1* locus with the *bar1*Δ::*hisG-URA3-hisG* allele. Replacement of *BAR1* with *hisG-URA3-hisG* was selected for after transformation by growth on -Ura medium and confirmed based on super sensitivity of the resulting strains to pheromone in halo assays and by PCR analysis using genomic DNA as template with primer pairs 967/968 and 966/972. The *bar1*Δ::*hisG* allele was generated from the resulting strains by selection on 5-fluororotic acid (5-FOA, 0.1% w/v) medium *(44)*. This medium provides a positive selection for isolates in which the *URA3* marker is excised by recombination within the direct *hisG* repeats *(45)*.

<u>*^P^FUS1-UBIY*Δ*KGFP^Sp^HIS5* and *^P^FUS1*(*PRE\**)-*UBIY*Δ*KGFP^Sp^HIS5* reporter gene strains BY4741-68 and BY4741-169</u> were derived from BY4741-66 by transformation with SacI-SalI digested pNC1146 or pNC1152, respectively and selection on −His medium. The integration of the reporter gene cassette in each strain was confirmed by PCR analysis using BY4741-68, and BY4741-169 genomic DNA as template with primer pair 867/1214.

<u>*far1*Δ_*0*_::*KanMX4 ^P^FUS1-UBIY*Δ*KGFP^Sp^HIS5* strain BY4741-130</u> was derived from BY4741-68 by transformation with a 3.3 kb fragment that was PCR amplified using BY4741-70 DNA as template with primer pair 1208/1209 and selection on G418 medium (200 *μ*g/ml). Replacement of the *FAR1* locus with the *far1*Δ_*0*_::*KanMX4* allele was confirmed by PCR analysis using BY4741-130 genomic DNA as template with primer pair 1210/881.

<u>*dig2*Δ_*0*_::*kanMX4 ^P^FUS1-UBIY*Δ*KGFP^Sp^HIS5* strain BY4741-110</u> was derived from BY4741-68 by transformation with a 1972 bp fragment that was PCR amplified using BY4741-29 genomic DNA as template with primer pair 1156/1157 and selection on G418 medium (200 *μ*g/ml). Replacement of the *DIG2* locus with the *dig2*Δ_*0*_::*kanMX4* allele was confirmed by PCR analysis using BY4741-110 genomic DNA as template with primer pair 868/881.

#### Strains constructed using the “Delitto Perfetto” approach (46)

<u>*^P^STE5-STE12 P_FUS1_-UBIY*Δ*KGFP^Sp^HIS5* strain BY4741-103</u> was derived from BY4741-68 by replacing the entire *SAM35-STE12* intergenic region (−1 to −485 from the *STE12* ATG codon) with 815 bp from the *ARX1-STE5* intergenic region (−1 to −815 bp from the *STE5* ATG codon). This promoter replacement was chosen because *STE5* is expressed constitutively at levels comparable to *STE12* basal expression (see below). The first step for construction of this strain was to replace a 94 bp region that encompasses the *STE12 UAS* with the PCR generated *ste12*Δ^*UAS*^::*CORE-UK* allele. pCORE-UK *(37)* was the template for the first round of PCR synthesis with primer pair 1019/1091. The resulting PCR product served as template for the second round with primer pair 1020/1093. Replacement of the *STE12 UAS* region with the CORE-UK cassette was selected for on -Ura medium and confirmed by PCR analysis using genomic DNA as template with primer pair 1194/881. In the second step, the *ste12*Δ^*UAS*^::*CORE-UK* allele was replaced with a *P_STE5_* fragment generated by three rounds of PCR. Genomic DNA was the template for the first round of PCR synthesis with primer pair 1121/1124. The PCR product from the previous round served as template for second and third round synthesis with primer pairs 1125/1127 and 1126/1128, respectively. The amplified 815 bp from the *STE5* intergenic sequence is flanked by primer derived sequences (61 bp on the 5’ end and 69 bp on the 3’ end) that target the PCR fragment to the *STE12* locus. The CORE-UK replacement was counter selected for on 5-FOA medium and verified by the concomitant reversion to G418 sensitivity. PCR analysis using BY4741-103 genomic DNA as template with primer pair 1194/1116 confirmed the integration at the *STE12* locus. DNA sequence analysis of the resulting PCR product with the amplifying primers confirmed the sequence fidelity of *P_STE5_* driving *STE12* expression.

#### Strains constructed using the PCR based gene deletion or modification method (47)

<u>*STE12-3xmyc*::*KanMX6 and P_STE5_-STE12-3xmyc::KanMX6* strains BY4741-105 and BY4741-132</u> were derived from BY4741-68 and BY4741-103, respectively by adding a C-terminal triple myc-tag to the *STE12* coding sequence. A cassette with the 3xmyc tag and the KanMX6 selectable marker was amplified using two rounds of PCR. In the first round of PCR pYM4 *(48)* was template DNA with primer pair 1119/1110. The second round PCR used the product of the first round as template with primer pair 1120/1111. Insertion of the tag was confirmed by PCR analysis using BY4741-105 and BY4741-132 genomic DNA as template with primer pair 903/881. DNA sequence analysis of the resulting PCR product with primers 903 and 1015 confirmed the sequence fidelity of the tag.

<u>*dig1*Δ_*0*_::*hyg* strains BY4741-147 and BY4741-148</u> were derived from BY4741-110 and BY4741-137, respectively by transformation with a 1890 bp PCR amplified *dig1*Δ_*0*_::*hyg* allele and selection on hygromycin B (200 *μ*g/ml) (Sigma Aldrich,) medium. The allele was amplified in three rounds of PCR. In the first round of PCR, pCORE-UH was template DNA with primer pair 1177/1179. The second round and third round of PCR used the product of the previous round as template with primer pair 1178/1180 and 1181/1182, respectively. Replacement of the *DIG1* locus with the *dig1*Δ_*0*_::*hyg* allele was confirmed by PCR analysis using BY4741-147 and BY4741-148 genomic DNA as template with primer pair 822/1148.

### Cell Extract Preparation and immunoblotting

The following procedure was performed to determine the phosphorylation state and relative amount of Fus3 and Kss1 in response to a pheromone stimulus. Cells either untreated or treated with 50 nM α-factor for different durations (as described in the figure legend) were harvested in TCA (5 % final concentration), washed with 10 mM NaN3 and pellets frozen at −80 °C. To prepare cell extracts, glass-bead lysis in TCA was performed as described before *(49)*. DC protein assay (Bio-Rad, Hercules, CA) was used to determine protein concentration. 25 *μ*g total protein was loaded per sample. Proteins were resolved on 10 % SDS-PAGE, transferred to nitrocellulose and detected by immunoblotting with Phospho-p44/42 MAPK antibodies at 1:500 (9101L, Cell Signaling Technology, Danvers, MA), Fus3 antibodies at 1:500 (sc-6773, Santa Cruz Biotechnology, Dallas, TX), and anti-G6PDH at 1:50,000 (A9521, Sigma-Aldrich, St. Louis, MO). Immunoreactive moieties were detected by chemifluorescent detection (Pierce ECL Plus, Thermo Fisher Scientific, Rockford, IL) of horseradish peroxidase-conjugated (HRP) antibodies (anti-rabbit, 170-5046, Bio-Rad, Hercules, CA; anti-goat, sc-2768, Santa Cruz Biotechnology, Dallas, TX; or anti mouse, A90-103P, Bethyl Laboratories, Montgomery, TX) at 1:10,000. Blots were scanned using Typhoon Trio+ (GE Healthcare, Little Chalfont, UK) and band intensity was quantified using Fiji (National Institute of Health).

The following procedure was performed to compare basal expression levels of Ste12 under either the native pheromone inducible promoter or a noninducible promoter, *^P^STE12* and *^P^STE5*, respectively. Cultures of BY4741-68 (*STE12*, untagged), BY4741-104 (*STE12-3xmyc*), and BY4741-133 (*^P^STE5-STE12-3xmyc*) were grown to a cell density of 1×10^7^ cells/mL in YPD, and 10 mL of each were harvested by centrifugation. The Ota protein extract protocol (Mattison et al., 1999) was followed to yield cell lysates that were then mixed in a 1:1 ratio with SDS running buffer and boiled. 10 *μ*L of each sample was run on an 8 % SDS-PAGE gel and transferred to nitrocellulose. The membrane was blocked for 1 hr at room temperature with 5 % milk in TBST. The membrane was then incubated with 1:1,000 of the primary goat anti-cMyc antibody (Bethyl Laboratories) in 5 % milk in TBST overnight at 4 °C. After washing in TBST, the membrane was incubated in 1:10,000 HRP-conjugated rabbit anti-goat secondary (Santa Cruz) in 10 mL of TBST with 200 *μ*L 5 % milk in TBST for 1 hr. The membrane was visualized using Western Lightning ECL Pro (PerkinElmer, Waltham, MA) and the ChemiDoc MP imaging system (Bio-Rad). The membrane was washed in TBST and incubated in 10 mL of stripping buffer at 65 °C for 45 min. The same protocol was then followed with 1:50,000 rabbit anti-G6PDH (A9521, Sigma-Aldrich) for the primary and 1:10,000 goat anti-rabbit HRP-labeled antibody (170-5046, Bio-Rad) for the secondary.

### Microfluidics

To generate time-dependent pheromone concentrations we used a microfluidics device and robotic automation which is part of the Dial-a-Wave system developed by the Laboratory of Dr. Jeff Hasty at USCD *(50)*. The device consists of a narrow chamber where cells are loaded and imaged, and two input ports; one containing pheromone and the other only containing media. When one of the input channels is positioned higher than the other, the fluid from that channel is at a higher pressure and flows into the chamber housing the cells. By alternating the height of the input channels, we can turn pheromone on and off in the chamber. The input channel also contained a 1:1,000 dilution of stock Alexa Fluor 647 (Invitrogen) dye, which has a similar diffusion coefficient as pheromone. The fluorescent signal is quantified in the chamber as the height of the input channels is alternated. Typically, it takes 2-5 min for the dye to equilibrate inside the chamber after switching the channels. This is much faster than the timescale of transcriptional response in the mating pathway. During the experiments, switching is automated using a step motor. Detailed microfluidics can be found in a methods review *(51)*.

### Microscopy

All experiments were performed in a microfluidic device using cell culturing methods as described in the supplement to Hao et al. (2008 Mol. Cell 30:649-656). Alexa 647 dye was added to pheromone containing media to enable imaging of the chamber and verification that dye (and by inference pheromone) turned on and off within less than 20 seconds. For experiments done at 50 nM constant pheromone and a single 200-minute pulse of 50 nM pheromone time-lapse microscopy was performed using a Nikon Ti-E inverted fluorescence microscope with Perfect Focus, coupled with Hamamatsu Orcaflash 4.0 digital camera and a Lumen Dynamics C-Cite LED light source system. Images were taken using a Nikon Plan Apo VC X60 oil immersion objective (NA 1.40 WD 0.17 MM). Images were taken every 5 minutes for pulses of stimulus and every 10 minutes for constant stimulus in the brightfield and green channels. Images were acquired in the far-red channel every other time point. The lowest LED intensity setting (5%) was used to prevent photobleaching and phototoxicity. Cells were imaged for 20 min prior to exposure to pheromone and for 10 hours thereafter

For all other experiments time-lapse microscopy was performed using a Nikon Ti-E 2000 inverted fluorescence microscope with Prior stage, coupled with Hamamatsu OrcaII Monochrome camera and a Prior Lumen200 light source system. Images were taken using a Nikon Plan Apo VC X60 oil immersion objective (NA 1.40 WD 0.17 MM).

### Image analysis

To improve the quality of segmentation images were edited in ImageJ by first subtracting the background, using the unsharp mask filer with a radius of 2.0 and a mask weight of 0.5, using the Gaussian blur filter with a radius of 3.0, and finally subtracting the background again. These images were then used to perform image segmentation using SchnitzCells *(52)*. The resulting segmentation was checked and corrected manually. The individual cells were tracked based on the position of each cell’s centroid and used to generate single cell traces of GFP fluorescence. All data is reported as the average of 90 or more single cell traces. For single pulses, analysis of wildtype and mutant strains was performed including and excluding daughter cells born after stimulus was removed. The comparison showed that excluding daughter cells did not change the average transcriptional response. We also compared transcriptional induction of cells in the G1, S and G2/M phases of the cell cycle when cells were exposed to 50 nM pheromone. This comparison revealed that cells in all phases of the cell cycle respond to pheromone by inducing transcription but those in S-phase respond more slowly than those in G1 and G2/M (**Fig. S7**).

### Model Development

Our experimental investigations revealed that the mechanisms responsible for persistent gene expression following removal of pheromone must occur downstream of MAP kinase activity. Therefore, we do not explicitly consider upstream signaling events and start our model at the level of the MAP kinase. The following equations govern the concentrations of inactive, [*MAPK*], and active, [*ppMAPK*], Fus3 and Kss1:

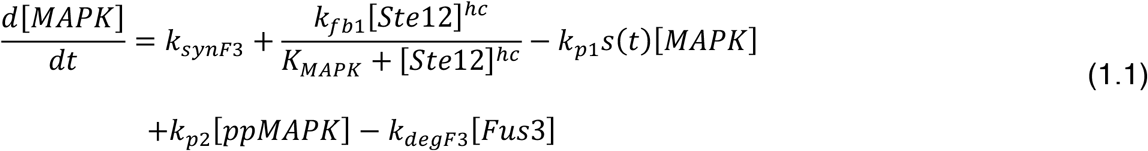

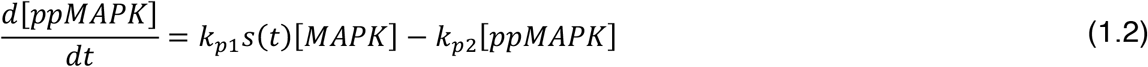

where *k*_*synF*3_ is the constitutive synthesis rate of Fus3, *k*_*degF*3_ is the MAPK degradation rate, *k*_*p*1_*s*(*t*) is the pheromone dependent activation rate of MAPK, and the term *k*_*fb*1_[*Ste*12]^*hc*^/(*K_MAPK_* + [*Ste*12]^*hc*^) models synthesis of Fus3 due to Ste12 dependent gene transcription.

Our model focuses on mechanisms that regulate Ste12-dependent gene expression. The regulatory mechanism we consider are self-induction of Ste12 (positive feedback), pheromone-dependent degradation of Ste12 (negative feed forward), and Ste12 inactivation by Dig1/2. For simplicity, we assume that the concentration of Dig1/2 remains constant, and when in a Ste12-Digs heterodimer Ste12 is protected from degradation. The equations that govern the concentration of free Ste12, [*Ste*12], and Ste12-Digs heterodimer, [*Ste*12_*Digs*_] are given by:

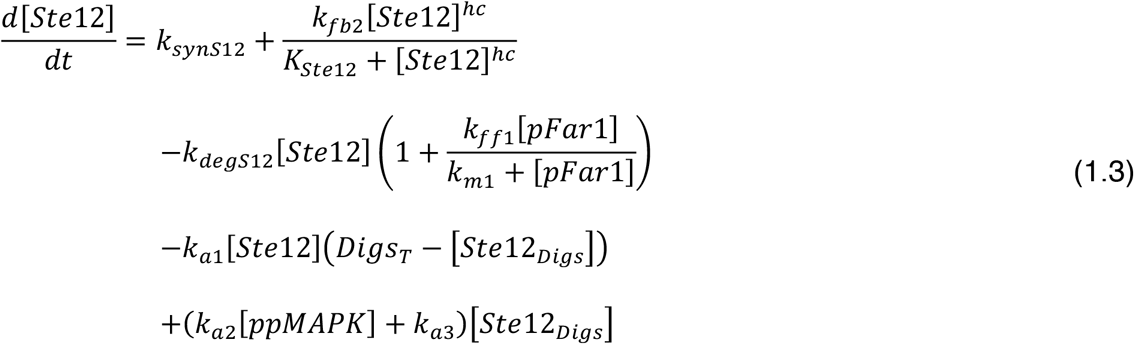

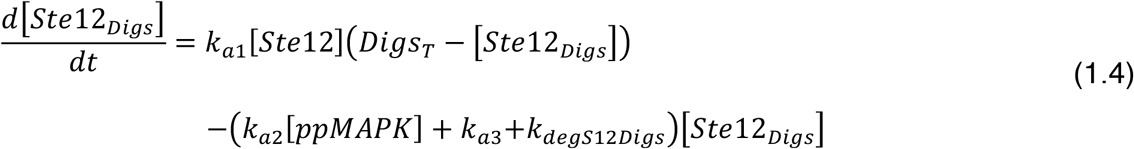

where *k*_*synS*12_ is the constitutive synthesis rate, *k*_*degS*12_ is the basal degradation rate of free Ste12, *k*_*degS*12*Digs*_ is the basal degradation rate of Ste12 in the Ste12-Digs complex, the term *k*_*ff*1_[*pFar*1]/*k*_*m*1_ + [*pFar*1] models the pheromone-dependent increase in the degradation rate, which depends on active Far1, and the term *k*_*fb*2_[*Ste*12]^*hc*^/*K*_*Ste*12_ + [*Ste*12]^*hc*^ model’s synthesis of Ste12 due to Ste12 dependent gene transcription. The terms in the third and fourth lines of Eq. (3) and in Eq. (4) represent the formation and dissociation of Ste12-Digs heterodimer. In this term, *Digs_T_* represents the total Dig1/2 concentration, which is assumed to remain constant, *k*_*a*1_ is the association rate constant, k_rsd2_ is the dissociation rate constant in the absence of pheromone and *k*_*a*2_[*ppMAPK*] is MAPK dependent increase in dissociation rate.

Since degradation of Ste12 is dependent on active Far1, our model includes Far1 dynamics. The equations that govern the concentrations of active Far1, [*pFar*1], and inactive Far1, [*Far*1] are given by:

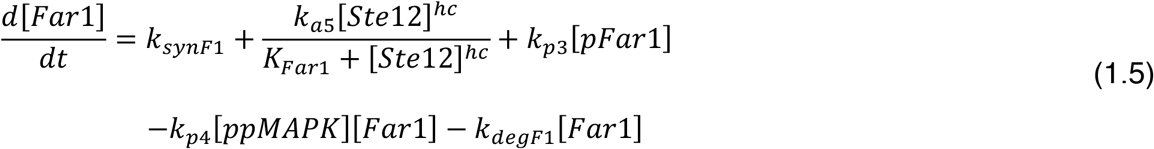

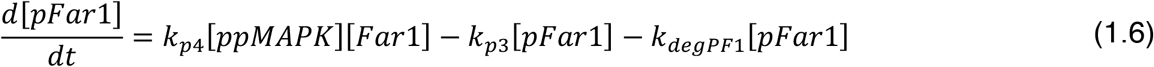

where *k*_*synF*1_ is the constitutive synthesis rate of Far1, *k*_*p*3_ is the dephosphorylation rate of active Far1, *k*_*p*4_[*ppFus*3] is the pheromone dependent rate of Far1 activation, *k*_*degF*1_ is the degradation rate of inactive Far1, *k*_*degPF*1_ is the degradation rate of active Far1, and the term *k*_*a*5_[*Ste*12]^*hc*^/*K*_*Far*1_ + [*Ste*12]^*hc*^ models synthesis of Far1 due to Ste12 dependent gene transcription.

The final component included in our model is GFP, which is expressed from a FUS1 promoter. GFP is included so we can directly compare our experimental data with the model. The equation that governs GFP, [*GFP*], synthesis and degradation are given by:

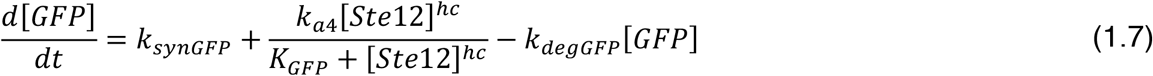

where *k_synGFP_* is the constitutive synthesis rate of GFP, *k_degGFP_* is the degradation rate of GFP, which is known as we used a short lived GFP with a well-established half-life, and the term *k*_*a*4_[*Ste*12]^*hc*^/*K_GFP_* + [*Ste*12]^*hc*^ models synthesis of GFP due to Ste12 dependent gene transcription.

### Modeling pheromone signal

In our model the pheromone signal comes in at the level of the MAPK, Fus3, which is activated in a pheromone dependent manner. We model upstream activation of the pathway as a piecewise linear function. We specify a slope *m_on_* that describes the rate at which signal activity increase following pheromone exposure and assume the signaling turns of instantaneously following removal of pheromone. The maximum input signal activity is k_p1_. For pulse trains of period p (on+off phase), the maximum input signal (*s_max_*) achieved during the simulation (*s_max_*) is:

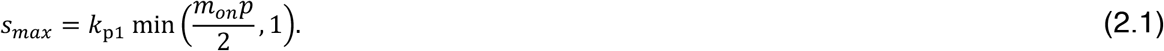

Then we use the following piecewise linear equation to describe the temporal signal profile for periodic stimulus:

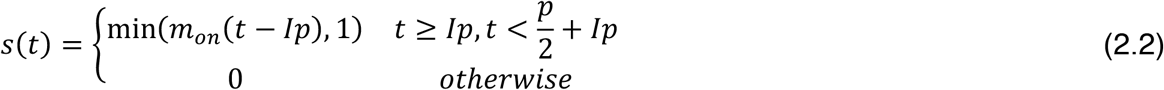

where *t* is the time and *I* is the pulse number defined by floor (*t/p*).

For single pulses of stimulus, the signal is described by the following equation:

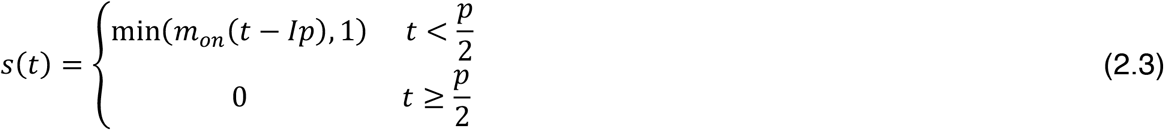

### Parameter estimation

To perform parameter estimation, we used an evolutionary algorithm. Evolutionary algorithms have the goal of optimizing a solution to a problem and are inspired by elements of biological evolution including recombination, mutation, and selection. In our application we aim to optimize the fit of our model to experimental data by finding parameter sets that minimize the error between the experimental and simulated data. We implemented the algorithm using DEAP (Distributed Evolutionary Algorithms in Python) which is a user friendly framework for building and executing evolutionary algorithms *(53)*.

The evolutionary algorithm is broken down into three main functions: simulation, scoring, and evolution. The algorithm is initiated by selecting parameter sets from specified uniform random distributions. In the simulation function, these parameters are used to simulate the model. In the scoring function, the difference between the simulation and the experimental data is the quantified using the mean absolute error. In the evolution function, the best parameters are chosen through a tournament. Then those parameters go through mating and mutation. Mating was simulated using a two-point crossover function and mutation was simulated using a polynomial bounded mutation function. The resulting parameter sets are then returned to the simulation function and the process starts over again. The algorithm continues to optimize parameter sets for a specified number of generations.

We performed all model fitting with 100 generations of 500 individuals because these numbers were typically sufficient for convergence of the score function. The mating and mutation functions were chosen because they worked best with synthetic data sets, specifically to optimize models with hill functions. Similarly, the hyperparameters (mutation rate, crossover rate, and tournament size) were selected based on their efficiency in fitting the synthetic data set best.

### Scaling experimental data

It was necessary to scale the experimental data sets to account for differences in experimental conditions, such as light source and intensity. We first normalized the wildtype time series for constant 50 nM α-factor to have a maximum of 1. Then all data sets generated using 50 nM α-factor were scaled to align with the wildtype response during the initial on phase of pheromone exposure. For example, the choice of scaling factor for a 45 min pulse experiment was based aligning the first 45 minutes of this times series with the response for the constant pheromone case.

For the pathway mutants, a series of constant stimulation experiments was done in quick succession using the same microscope settings. In this experiment data were collected for wildtype, *far1Δ, dig1Δdig2Δ*, PRE*-GFP, and *^P^STE5-STE12* strains. The wildtype data was multiplied by a scaling factor to align with the data used to train the model. The mutant responses were then multiplied by this scaling factor to correctly adjust their starting and maximal values. The response of the each of the mutants to a 90-miunte pulse of stimulus was then multiplied by a strain specific scaling factor to match the corresponding scaled constant response.

### Prediction response to low pheromone dose

To predict the low dose data set, we scaled the signal input signal on rate (*m_on_*) by 0.3 based on an estimation of the difference in the slopes for the temporal response to 50 nM and 10 nM constant pheromone. Using the parameter sets corresponding to the best fits to the 50 nM data but changing *m_on_*, we successfully simulated the 10 nM data (**Fig. 8, blue curves**).

### Predicting response of pathway perturbations

The model was used to predict the response of four mutations to the pathway, *far1Δ, dig1Δdig2Δ*, PRE*-GFP, and pSTE5-STE12. The *far1Δ* was described in the model by setting all parameters related to Far1 expression, degradation, or activation (*k*_*synF*1_,*k*_*a*5_,*k*_*p*3_,*k*_*p*4_*k*_*degF*1_,*k*_*degPF*1_) equal to zero. The *dig1Δdig2Δ* was described in the model by setting the total amount of Dig1 and Dig2 (*Digs_T_*) equal to zero. The PRE*-GFP was described in the model by setting the apparent dissociation constant for Ste12 binding to the pheromone responsive element (PRE) of the GFP promoter (*K_GFP_*) equal to 3.33 times the value given from the best fits corresponding to the reported the relative competition strength of 0.3 *(19)*. Finally, the pSTE5-STE12 was described in the model by setting the parameter responsible for Ste12 inducing its own transcription (*k*_*fb*2_) equal to zero.

## Author contributions

A.E.P., J.R.H., M.I.P., T.C.E. and B.E. developed the research strategy. A.E.P., M.I.P., J.R.H., T.C.E., and B.E. planned the experiments. A.E.P., M.I.P., J.R.H., G.D., and B.E. performed the experiments. A.E.P., M.I.P., J.R.H., T.C.E., and B.E analyzed the experimental data. A.E.P., J.R.H., B.E. and T.C.E. developed the mathematical model. A.E.P. performed the computational analysis. A.E.P., H.G.D., T.C.E., and B.E. wrote the manuscript with input from the other authors.

## Data and materials availability

All data needed to evaluate the conclusions in the paper are present in the paper or the Supplementary Materials. All code used for analysis is available on GitHub. All materials are available upon request from the corresponding authors.

