## Supplement for "A predictive model of gene expression reveals the role of regulatory motifs in the mating response of yeast"

**Simple linear model**

To determine if persistent gene expression following removal of pheromone could be explained by a lag between signal input and transcriptional induction, we used a simple linear model (**Fig. S1**). It has four species, $X_1$, $X_2$, $X_3$, and $X_4$, where $X_4$ represents the transcriptional activation quantified by GFP fluorescence. Four species were chosen since three steps were required to capture the dynamics of the response to stimulus. The following equations govern the behavior of the simple model:

$$\frac{d[X_1]}{dt} = s - d_1 X_1 \quad (1.1)$$

$$\frac{d[X_2]}{dt} = k_1 X_1 - d_2 X_2 \quad (1.2)$$

$$\frac{d[X_3]}{dt} = k_2 X_2 - d_3 X_3 \quad (1.3)$$

$$\frac{d[X_4]}{dt} = k_3 X_3 - d_4 X_4 \quad (1.4)$$

where $k_1$, $k_2$, and $k_3$are the rates at which $X_1$, $X_2$, and $X_3$ generate $X_2$, $X_3$, and $X_4$ respectively, $d_1$, $d_2$, $d_3$, and $d_4$ are the rates at which $X_1$, $X_2$, $X_3$, and $X_4$ are degraded, and $s$ is the signal, which is modeled as is described in equation 1.10 in the main text.

This model was then fit to the “on phase” single pulse data (only those timepoints when stimulus was present) using least squares regression. Since the initial value of $X_4$ is zero for this system of equations, we added 0.14 to the solution so that the simulations would align with the initial values of the experimental data.

**Determining parameter ranges**

***Degradation rates***

The ranges for the basal degradation rates Fus3, Ste12, and Far1 ($k_{degF3}, k_{degS12}, k_{degS12D}, k_{degF1}, k_{degPF1}$) were determined based on the reported half-lives of the three proteins *(1)*. These parameters were allowed to range from rates corresponding to 10 times higher to 10 times lower than the reported half-lives. The degradation rate of GFP ($k_{degGFP}$) was set to 0.01 based on the known half-life of the short lived GFP.

***Synthesis rates***

The synthesis rates for Fus3, Ste12 and Far1 ($k_{synF3}, k_{synS12}, k_{synF1}$) were determined to keep the basal abundance of each molecule per cell between reported ranges (Fus3: 1,000-10,000, Ste12: 2,000-6,000, Far1: 40-2,000) *(2–17)*. The values for the synthesis rate of GFP ($k_{synGFP}$) ranged between the smallest and largest allowable rates for Fus3, Ste12 and Far 1 synthesis.

Transcriptional induction by Ste12 was modeled using Hill equations. The $V_{max}$ values ($k_{fb1}, k_{fb2}, k_{a5}, k_{a4}$) for each protein were set to allow a range of 1 to 1,000 additional molecules to be produced per minute for Ste12, Fus3 and Far1 and an extra 1 to 50,000 molecules for GFP. All Hill constants were taken to have a value of 2. The range of values for the constants ($K_{MAPK}, K_{Ste12}, K_{Far1}, K_{GFP}$) that determine the Ste12 concentration at which the Hill function is at its half maximum value was 0.24 μM to 476.2 μM. Assuming a cell volume of $42 \mu m^3$, this range corresponds to a range of Ste12 molecules of 1 to 20,000.

***Reaction rates***

For second order reactions, the range for the association rate constants ($k_{a2}, k_{a1}, k_{p4}$) were set so that the maximum rate was the diffusion limit, calculated using $k = 4\pi Dr$ where $D = 15\mu m^2/s$ is the combined diffusivity of the two reactants and $r = 0.01\mu m$ is the reaction radius. The minimum rate was set so the time scale for one reaction to occur was no longer than 1 hour. The basal dissociation rate for $Ste12_{Digs}$ ($k_{a3}$) was set so there would be an amount of $Ste12_{Digs}$ that was in the range of the total Dig concentration $Digs_T$.

### *Phosphorylation and dephosphorylation rates*

For all phosphorylation and dephosphorylation reactions,  rates are proportional to the concentration of substrate being phosphorylated or dephosphorylated ($k_{p1}, k_{p2}, k_{p3}$), we assigned a range of $10^{-5}$-$10^{-1}$ $s^{-1}$ based on previous work *(18)*.

### *Abundance of Digs*

The amount of Dig1 and Dig2 in the cell ($Digs_T$) was allowed to range from 10 times lower than the lowest reported abundance of either Dig to 10 times higher than the highest reported abundance of either Dig. Micromolar concentrations were calculated assuming a nuclear volume of 2.91µm$^3$.

### *Input Signal*

The input signal was assumed to piecewise linear.  The slope at which the signal increased ($m_{on}$) was allowed to range between 45 and 250 minutes to reach the maximum of 1. The signal was assumed to turn off immediately following removal of pheromone.

| Rate constant | Range allowed | Reaction | Reaction rate |
|---|---|---|---|
| $k_{synF3}$ | $10^{-6.9} - 10^{-1.6}$ | $\emptyset \rightarrow Fus3$ | $k_{synF3} + \frac{k_{fb1}[Ste12]^{hc}}{K_{Fus3} + [Ste12]^{hc}}$ |
| $k_{fb1}$ | $10^{-10.3} - 10^{-1.7}$ | $\emptyset \rightarrow Fus3$ | $k_{synF3} + \frac{k_{fb1}[Ste12]^{hc}}{K_{Fus3} + [Ste12]^{hc}}$ |
| $K_{Fus3}$ | $10^{-4.5} - 10^{0.0}$ | $\emptyset \rightarrow Fus3$ | $k_{synF3} + \frac{k_{fb1}[Ste12]^{hc}}{K_{Fus3} + [Ste12]^{hc}}$ |
| $k_{p1}$ | $10^{-5.0} - 10^{-1.0}$ | $Fus3 \rightarrow ppFus3$ | $k_{p1}s(t)[Fus3]$ |
| $k_{p2}$ | $10^{-5.0} - 10^{-1.0}$ | $ppFus3 \rightarrow Fus3$ | $k_{p2}[ppFus3]$ |
| $k_{degF3}$ | $10^{-3.6} - 10^{-1.5}$ | $Fus3 \rightarrow \emptyset$ | $k_{degF3}[Fus3]$ |
| $k_{synS12}$ | $10^{-7.1} - 10^{-1.6}$ | $\emptyset \rightarrow Ste12$ | $k_{synS12}$ |
| $k_{fb2}$ | $10^{-10.5} - 10^{-1.7}$ | $\emptyset \rightarrow Ste12$ | $\frac{k_{fb2}[Ste12]^{hc}}{K_{Ste12} + [Ste12]^{hc}}$ |
| $K_{Ste12}$ | $10^{-4.5} - 10^{0.0}$ | $\emptyset \rightarrow Ste12$ | $\frac{k_{fb2}[Ste12]^{hc}}{K_{Ste12} + [Ste12]^{hc}}$ |
| $k_{degS12}$ | $10^{-3.7} - 10^{-1.5}$ | $Ste12 \rightarrow \emptyset$ | $k_{degS12}[Ste12]\left(1 + \frac{k_{ff1}[pFar1]}{k_{m1} + [pFar1]}\right)$ |
| $k_{ff1}$ | $10^{-12.0} - 10^{6.8}$ | $Ste12 \rightarrow \emptyset$ | $k_{degS12}[Ste12]\left(1 + \frac{k_{ff1}[pFar1]}{k_{m1} + [pFar1]}\right)$ |
| $k_{m1}$ | $10^{-2.3} - 10^{0.0}$ | $Ste12 \rightarrow \emptyset$ | $k_{degS12}[Ste12]\left(1 + \frac{k_{ff1}[pFar1]}{k_{m1} + [pFar1]}\right)$ |
| $k_{a1}$ | $10^{-11.0} - 10^{4.0}$ | $Ste12 \rightarrow Ste12_{Digs}$ | $k_{a1}[Ste12](Digs_T - [Ste12_{Digs}])$ |
| $Digs_T$ | $10^{-2.8} - 10^{1.0}$ | $Ste12 \rightarrow Ste12_{Digs}$ | $k_{a1}[Ste12](Digs_T - [Ste12_{Digs}])$ |
| $k_{a2}$ | $10^{-11.0} - 10^{4.0}$ | $Ste12_{Digs} \rightarrow Ste12$ | $(k_{a2}[ppFus3] + k_{a3})[Ste12_{Digs}]$ |
| $k_{a3}$ | $10^{-12.0} - 10^{6.8}$ | $Ste12_{Digs} \rightarrow Ste12$ | $(k_{a2}[ppFus3] + k_{a3})[Ste12_{Digs}]$ |
| $k_{synF1}$ | $10^{-6.1} - 10^{-0.4}$ | $\emptyset \rightarrow Far1$ | $k_{synF1} + \frac{k_{a5}[Ste12]^{hc}}{K_{Far1} + [Ste12]^{hc}}$ |
| $k_{a5}$ | $10^{-9.5} - 10^{-0.7}$ | $\emptyset \rightarrow Far1$ | $k_{synF1} + \frac{k_{a5}[Ste12]^{hc}}{K_{Far1} + [Ste12]^{hc}}$ |
| $K_{Far1}$ | $10^{-4.5} - 10^{0.0}$ | $\emptyset \rightarrow Far1$ | $k_{synF1} + \frac{k_{a5}[Ste12]^{hc}}{K_{Far1} + [Ste12]^{hc}}$ |
| $k_{p3}$ | $10^{-5.0} - 10^{-1.0}$ | $pFar1 \rightarrow Far1$ | $k_{p3}[pFar1]$ |

| | | | |
|---|---|---|---|
| $k_{p4}$ | $10^{-11.0} - 10^{0.0}$ | $Far1 \rightarrow pFar1$ | $k_{p4}[ppFus3][Far1]$ |
| $k_{degF1}$ | $10^{-2.7} - 10^{-0.5}$ | $Far1 \rightarrow \emptyset$ | $k_{degF1}[Far1]$ |
| $k_{degPF1}$ | $10^{-3.7} - 10^{-0.5}$ | $pFar1 \rightarrow \emptyset$ | $k_{degPF1}[pFar1]$ |
| $k_{synGFP}$ | $10^{-7.1} - 10^{-0.6}$ | $\emptyset \rightarrow GFP$ | $k_{synGFP} + \frac{k_{a4}[Ste12]^{hc}}{K_{GFP} + [Ste12]^{hc}}$ |
| $k_{a4}$ | $10^{-10.5} - 10^{0.0}$ | $\emptyset \rightarrow GFP$ | $k_{synGFP} + \frac{k_{a4}[Ste12]^{hc}}{K_{GFP} + [Ste12]^{hc}}$ |
| $K_{GFP}$ | $10^{-4.5} - 10^{0.0}$ | $\emptyset \rightarrow GFP$ | $k_{synGFP} + \frac{k_{a4}[Ste12]^{hc}}{K_{GFP} + [Ste12]^{hc}}$ |
| $k_{degGFP}$ | $10^{-1.0}$ | $GFP \rightarrow \emptyset$ | $k_{degGFP}[GFP]$ |
| $hc$ | 2 | $\emptyset \rightarrow Fus3$<br>$\emptyset \rightarrow Ste12$<br>$\emptyset \rightarrow Far1$<br>$\emptyset \rightarrow GFP$ | $\frac{k_{fb1}[Ste12]^{hc}}{K_{Fus3}+[Ste12]^{hc}}, \frac{k_{fb2}[Ste12]^{hc}}{K_{Ste12}+[Ste12]^{hc}},$<br>$\frac{k_{a5}[Ste12]^{hc}}{K_{Far1}+[Ste12]^{hc}}, \frac{k_{a4}[Ste12]^{hc}}{K_{GFP}+[Ste12]^{hc}}$ |
| $m_{on}$ | $10^{-2.6} - 10^{-1.6}$ | | |

**Table S1. Parameter ranges for evolutionary algorithm.** This table includes all of the parameters in our model and the range used in the evolutionary algorithm for parameter estimation.

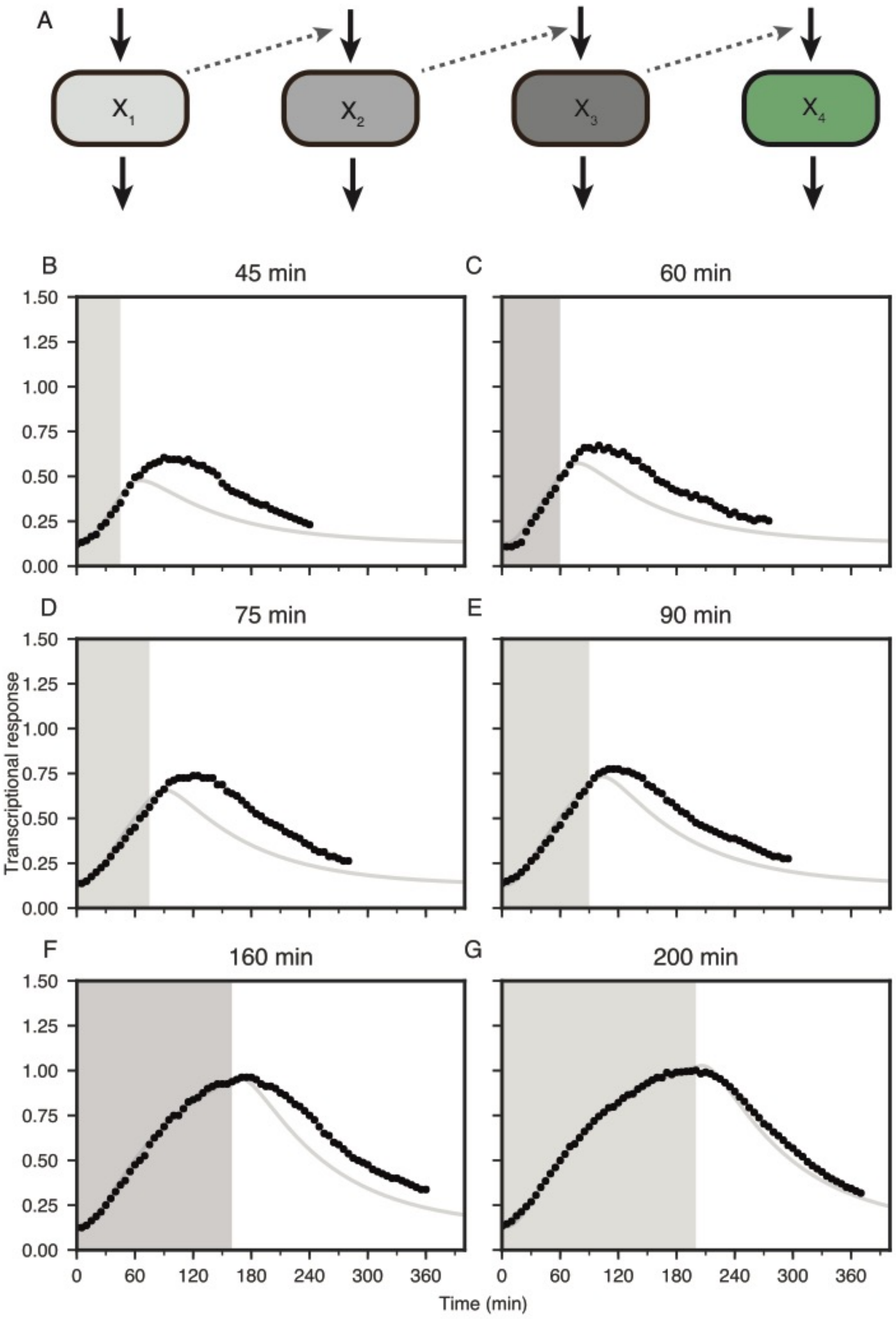


**Figure S1. Simple delay model.** (A) Schematic of a simple delay model used to determine whether the persistence could be attributed to a delay in transcription and translation. (B-G) Best fit (gray line) to the activation of the pathway (area shaded in gray) as found by least squares regression for $X_4$ in the simple delay model to the transcriptional response of wildtype strain (BY4741-68) to single pulses of stimulus (circles) for six different durations (45, 60, 75, 90, 160, and 200 min).

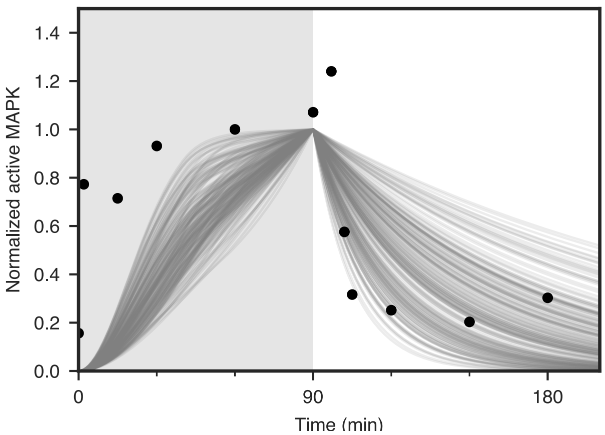


**Figure S2. MAPK activation dynamics.** Model simulations generated using the top 10% of parameters found by the evolutionary algorithm (gray lines) compared to the experimental data (circles) for wildtype strain (BY4741-68) MAPK response to a 90-min pulse of 50 nM pheromone. Gray shading indicates when mating pheromone is present in the time course. Note that in the model we do not distinguish nuclear from cytoplasmic MAPK, and, therefore, all active MAPK is available to drive transcription. This simplification may explain why the model underestimates total MAPK activity at early time points, because in cells only a fraction of active MAPK is in the nucleus.

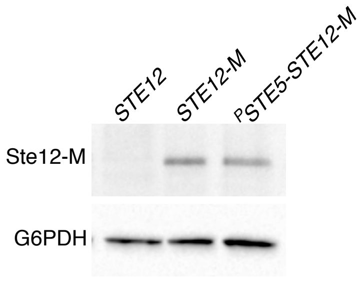


**Figure S3. *PSTE5-STE12* and wildtype have similar basal Ste12 abundance.** Ste12 abundance in *STE12*-3x*myc*, and *PSTE5-STE12*-3x*myc* strains were visualized using anti-cMyc antibodies. The *STE12* strain serves a negative control for detection of the Ste12-3xmyc fusion protein. Detection of G6PDH with anti-G6PDH antibodies provides as a loading control.

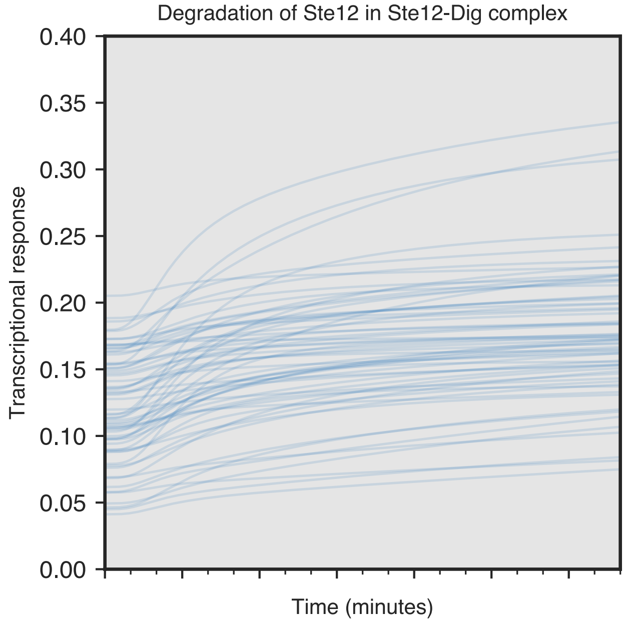


**Figure S4. Degradation of Ste12 in the Ste12-Dig complex prevents adaptation.** Parameter sets that did not adapt in simulations in the absence of Far1 were used to simulate the transcriptional response when binding of transcriptional repressors (Dig1 and Dig2) does not protect Ste12 from degradation (blue lines). This condition is simulated by Ste12 in complex having the same degradation rate as free Ste12. Simulations for elimination protective binding by Digs show no long-term adaptation.

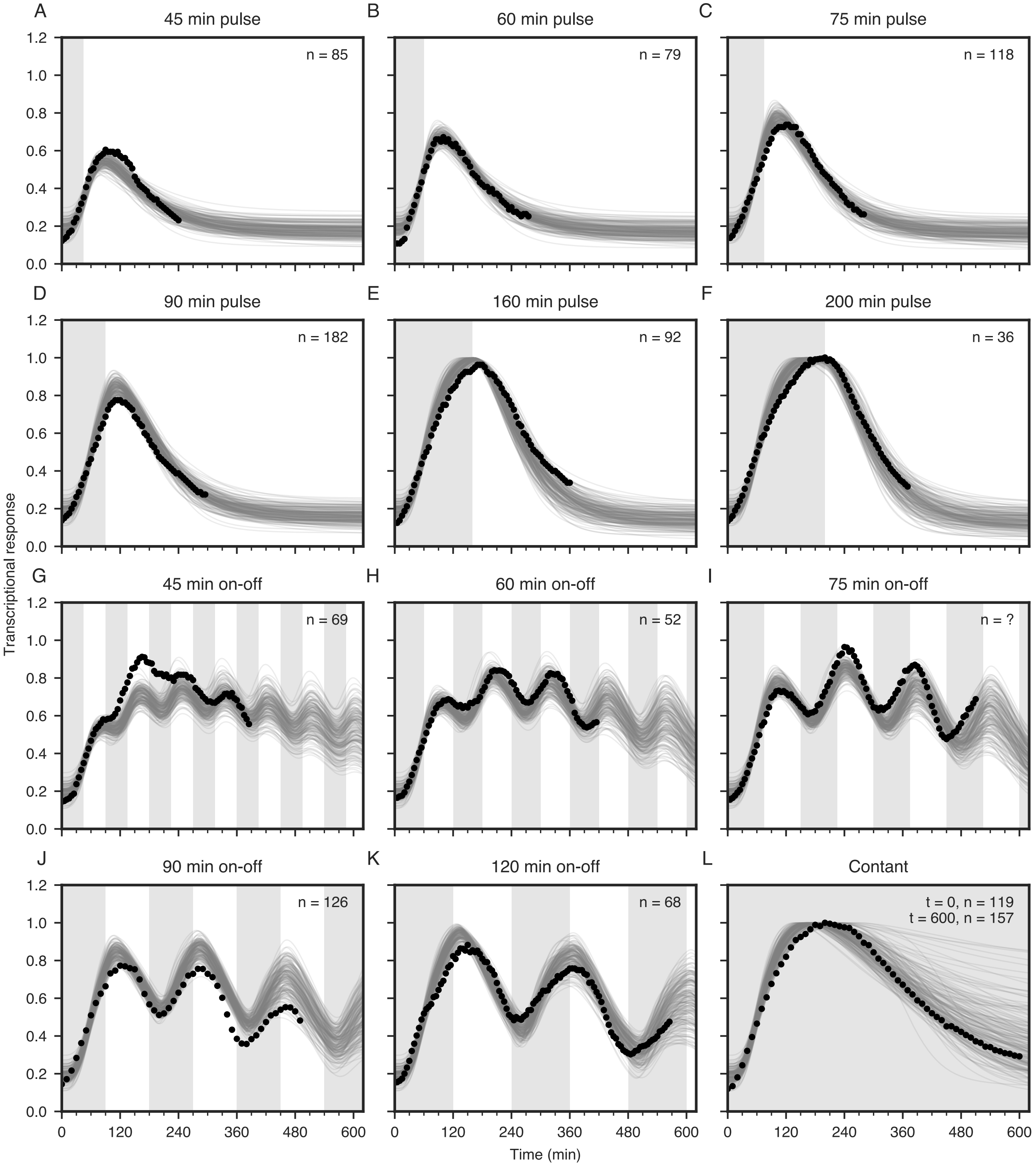


**Figure S5. Model captures response to dynamic stimulation.** Model simulations generated using the top 10% of parameters found by the evolutionary algorithm fit to training data from wildtype (constant, single pulse, and periodic stimulus) (BY4741-68), *far1Δ* (BY4741-130) (constant stimulus), and *$^{P}$STE5-STE12* (BY4741-103) (constant stimulus) strains (gray lines) compared to the experimental data (circles) for the wildtype strain (BY4741-68) transcriptional response to (A-F) six different pulse durations (45, 60, 75, 90, 160, and 200 min), (G-K) five different oscillatory stimulation profiles (45, 60, 75, 90, and 120 min on-off), and (L) constant stimulus. Gray shading indicates when mating pheromone is present.

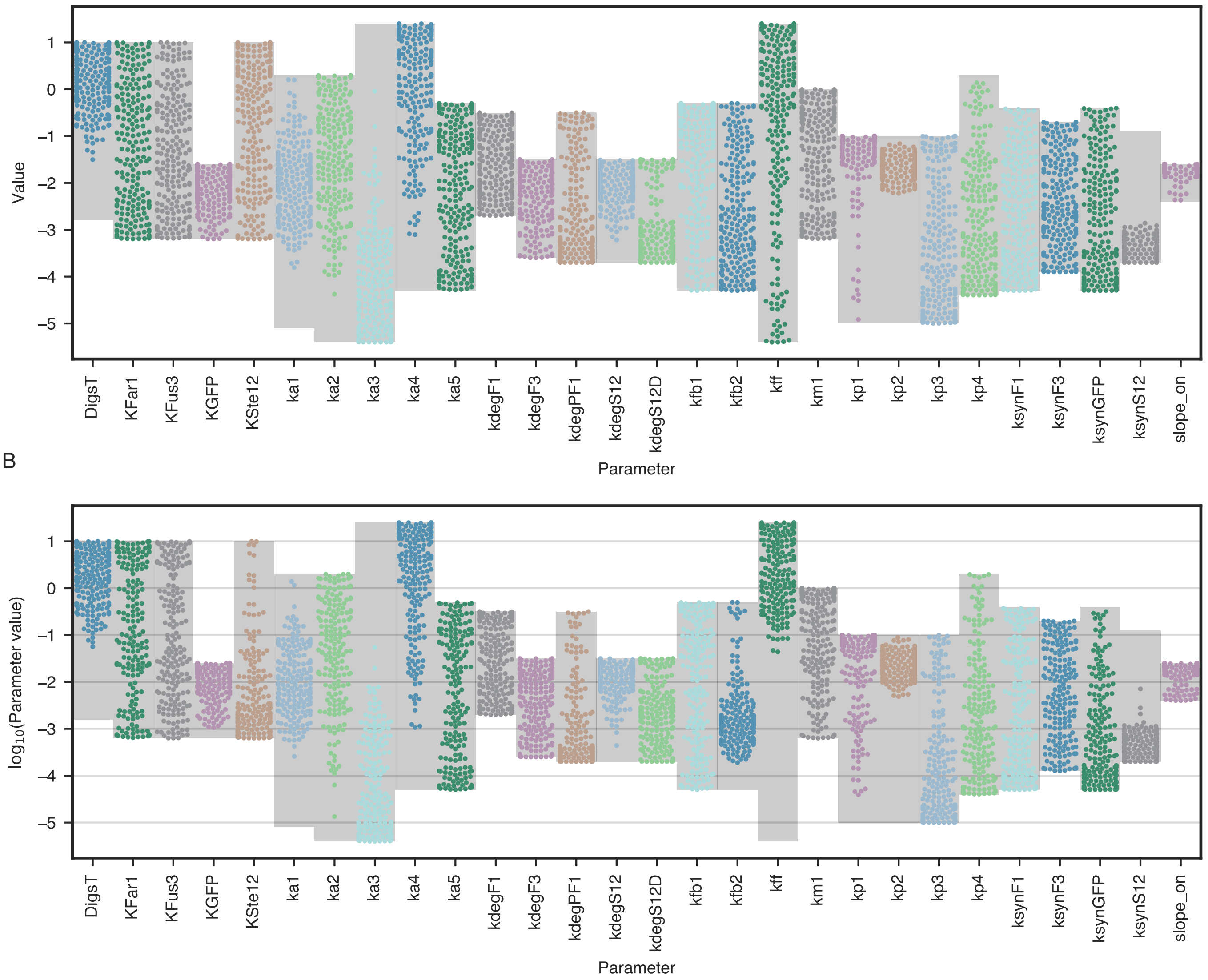


**Figure S6. Parameter distributions.** Distributions of parameters for the top 10% of model when fit to (A) wildtype only data (see Fig. 4 for fits) and (B) wildtype, *far1Δ*, and *$^{P}$STE5-STE12* data (see **Figs. S5** and **6** for fits). Gray boxes represent the allowed ranges for each parameter (see supplemental material for details on how range was chosen).

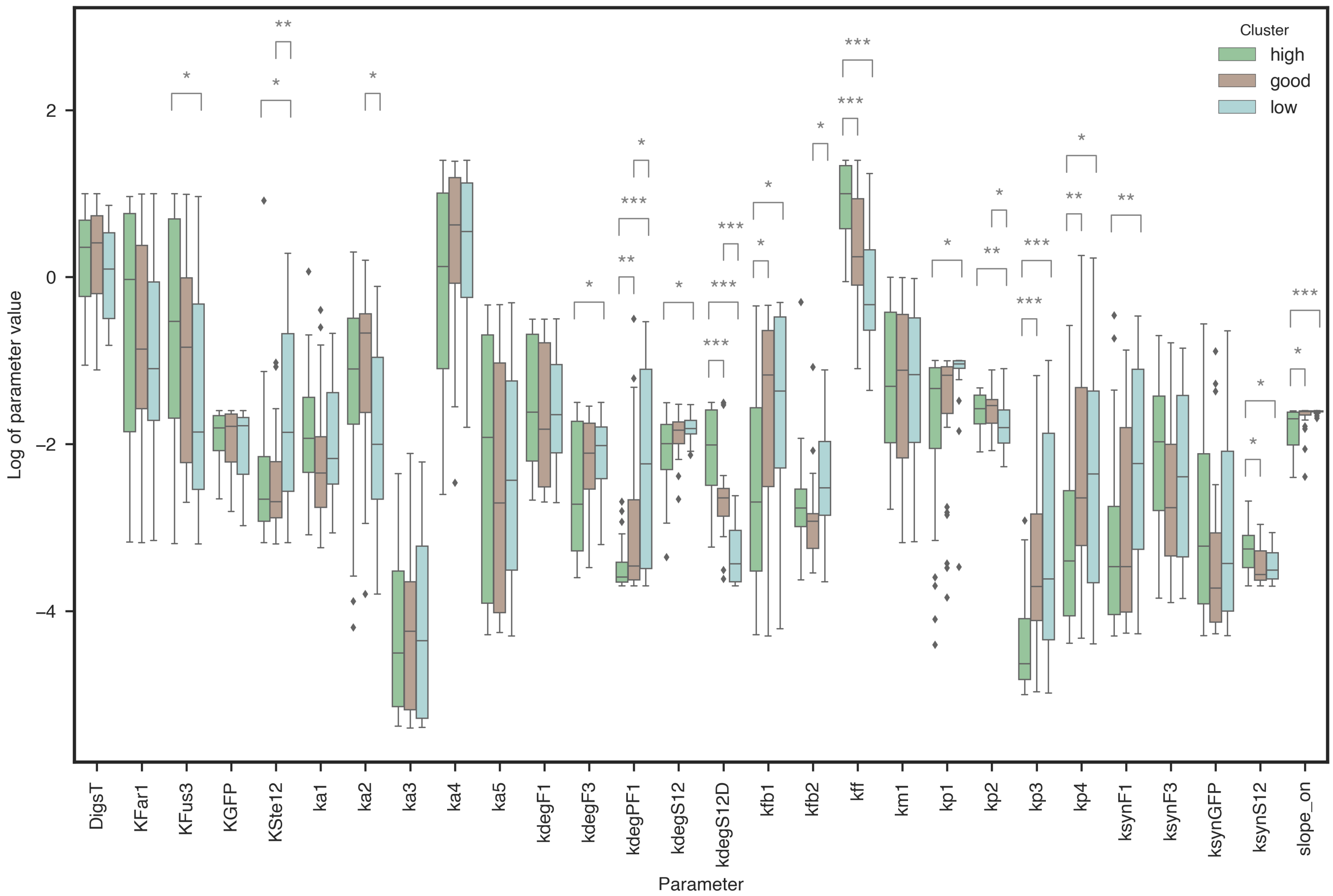


**Figure S7. Effect of parameter distributions on predicted response for *dig1Δdig2Δ*.** Analysis of parameter distributions within each of the clusters for *dig1Δdig2Δ response* (Fig 7C) for all estimated parameters (see Supplemental Table 1 for parameter descriptions). Significance values (*$p < 0.5$, **$p < 0.1$, and ***$p < 0.01$) were calculated using a t-test with a Bonferroni correction for multiple hypothesis testing.

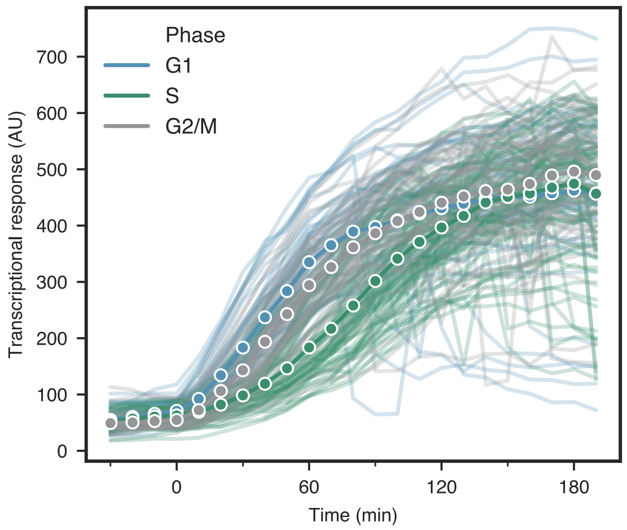


**Figure S8. Cell cycle dependence of transcriptional response.** Single cell and mean responses to constant stimulus for cells in G1 (blue), S (green), and G2/M (gray) phase of the cell cycle at the time stimulus is added. Cells in S phase at the time stimulus was administered are slower to respond but reach the same maximal transcriptional response.
